# On the state of protein function prediction: a report on the fourth CAFA challenge

**DOI:** 10.64898/2026.05.06.722942

**Authors:** Rashika Ramola, M. Clara De Paolis Kaluza, Damiano Piovesan, Yisu Peng, Parnal Joshi, Mahta Mehdiabadi, Federica Quaglia, Rita Pancsa, Lucía B. Chemes, Hongryul Ahn, Adrian M. Altenhoff, Ehsaneddin Asgari, Maria Cristina Aspromonte, Volkan Atalay, Giulia Babbi, Davide Baldazzi, Meet M. Barot, Asa Ben-Hur, Alfredo Benso, Daniel Berenberg, Jari Björne, Florian Boecker, Paolo Boldi, Joseph Bonello, Nicola Bordin, Piyush Borole, Ali Ebrahimpour Boroojeny, Renzhi Cao, Stefano Di Carlo, Rita Casadio, Elena Casiraghi, Jia-Ming Chang, Chen Chen, Tse-Ming Chen, Jianlin Cheng, Ssu Chiu, Alperen Dalkıran, Radoslav S. Davidović, Christophe Dessimoz, Rucheng Diao, Warith Eddine Djeddi, Tunca Dogan, Sean T. Flannery, Paolo Fontana, Marco Frasca, Lydia Freddolino, Branislava Gemović, Jesse Gillis, Filip Ginter, Vladimir Gligorijevic, Giuliano Grossi, Michael Heinzinger, Kyle Hippe, Robert Hoehndorf, Liisa Holm, Jie Hou, John R. Hover, Yen-Ting Huang, Emilio Ispano, Suraiya Jabin, Aashish Jain, David T. Jones, Suwisa Kaewphan, Yuki Kagaya, Jenna Kanerva, Daisuke Kihara, Maxat Kulmanov, Sunil Kumar, Lukasz Kurgan, Enrico Lavezzo, Jon Lees, Wen-Hung Liao, Han Lin, Michal Linial, Maria Littmann, Lizhi Liu, Tong Liu, Yi Wei Liu, Stavros Makrodimitris, Laura Manuto, Pier Luigi Martelli, Alice Carolyn Mchardy, Gabriela A. Merino, Diego H. Milone, Sarthak Mishra, Mohammad R. K. Mofrad, David Moi, Tsukasa Nakamura, Vijay Kumar Narsapuram, Maria Victoria Nugnes, Takeshi Obayashi, Dan Ofer, Alberto Paccanaro, Vladimir R. Perovic, Alessandro Petrini, Gianfranco Politano, Daniele Raimondi, Nadav Rappoport, Hafeez Ur Rehman, Maarten J. M. F. Reijnders, Marcel J. T. Reinders, P. Douglas Renfrew, Ahmet S. Rifaioglu, Alfonso E. Romero, Abhiman Saraswathi, Castrense Savojardo, Harry M. Scholes, Heiko Schoof, Yang Shen, Ian Sillitoe, Georgina Stegmayer, Amos Stern, Henri Tiittanen, Sumyyah Toonsi, Stefano Toppo, Petri Toronen, Mateo Torres, Gabriella Trucco, Giorgio Valentini, Nevena Veljkovic, Alex Warwick Vesztrocy, Vedrana Vidulin, Amelia Villegas-Morcillo, Antti Virtanen, Wim Vranken, Slobodan Vucetic, Cen Wan, Zheng Wang, Mark N. Wass, Robert M. Waterhouse, Sadok Ben Yahia, Haixuan Yang, Shuwei Yao, Ronghui You, Jeffrey Yunes, Chengxin Zhang, Yang Zhang, Chenguang Zhao, Xiaogen Zhou, Yi-Heng Zhu, Shanfeng Zhu, Hao Zhu, Gökhan Özsari, Burkhard Rost, Christine Orengo, Marc Robinson-Rechavi, Dannie Durand, Steven E. Brenner, Casey S. Greene, Sean D. Mooney, Silvio C. E. Tosatto, Iddo Friedberg, Predrag Radivojac

**Affiliations:** Khoury College of Computer Sciences, Northeastern University, Boston, MA, USA; Department of Biomedical Sciences, University of Padova, Padova, Italy; Bioinformatics and Computational Biology Program, Iowa State University, Ames, IA, USA; Department of Veterinary Microbiology & Preventive Medicine, Iowa State University, Ames, IA, USA; HUN-REN Research Centre for Natural Sciences, Institute of Molecular Life Sciences, Budapest, Hungary; Instituto de Investigaciones Biotecnológicas, Consejo Nacional de Investigaciones Científicas y Técnicas (CONICET), Escuela de Bio y Nanotecnologías (EByN), Universidad Nacional de San Martín, Buenos Aires, Argentina; Division of Data Science, The University of Suwon, Gyeonggido, South Korea; Comparative Genomics, SIB Swiss Institute of Bioinformatics, Lausanne, Switzerland; Department of Computer Science, ETH Zurich, Zurich, Switzerland; Qatar Computing Research Institute, HBKU, Doha, Qatar; Department of Computer Engineering, Middle East Technical University, Ankara, Turkey; Bologna Biocomputing Group, University of Bologna, Bologna, Italy; Oncogenetics and Functional Oncogenomics, CRO Aviano, National Cancer Institute, IRCCS, Aviano, Italy; Center for Data Science, New York University, New York, NY, USA; Department of Computer Science, Colorado State University, Fort Collins, CO, USA; Department of Control and Computer Engineering, Politecnico di Torino, Turin, Italy; Department of Computer Science, New York University, New York, NY, USA; Department of Computing, University of Turku, Turku, Finland; INRES Crop Bioinformatics, University of Bonn, Bonn, Germany; Dipartimento di Informatica Giovanni Degli Antoni, University of Milan, Milano, Italy; Computer Information Systems, University of Malta, Msida, Malta; Structural and Molecular Biology, University College London, London, England; Institute of Structural and Molecular Biology, University College London, London, UK; College of Science and Technology, Temple University, Philadelphia, PA, USA; Department of Computer Science, University of Illinois at Urbana-Champaign, Urbana, IL, USA; Department of Computer Science, Pacific Lutheran University, Tacoma, WA, USA; Department of Computer Science, University of Milan, Milano, Italy; Environmental Genomics and Systems Biology Division, Lawrence Berkeley National Laboratory, Berkeley, CA, USA; Department of Computer Science, National Chengchi University, Taipei City, Taiwan; Department of Electrical Engineering and Computer Science, University of Missouri Columbia, Columbia, MO, USA; NextGen Precision Health, University of Missouri Columbia, Columbia, MO, USA; Laboratory for Bioinformatics and Computational Chemistry, Institute of Nuclear Sciences Vinca, National Institute of the Republic of Serbia, University of Belgrade, Belgrade, Serbia; Department of Computational Biology, University of Lausanne, Lausanne, Switzerland; Department of Computational Medicine and Bioinformatics, University of Michigan, Ann Arbor, MI, USA; FST Manar, University of Tunis El Manar, Tunis, Tunisia; University of Jendouba, Jendouba, Tunisia; Department of Computer Engineering, Hacettepe University, Ankara, Turkey; Department of Bioinformatics, Hacettepe University, Ankara, Turkey; Department of Computer Science, Purdue University, West Lafayette, IN, USA; Research and Innovation Centre, Fondazione Edmund Mach, San Michele all’Adige, Italy; Department of Biological Chemistry, University of Michigan, Ann Arbor, MI, USA; Department of Physiology, University of Toronto, Toronto, ON, Canada; Donnelly Centre, University of Toronto, Toronto, ON, Canada; Cold Spring Harbor Laboratory, Cold Spring Harbor, NY; Prescient Design, Genentech, New York, NY, USA; School of Computation, Information and Technology (CIT), Technical University of Munich, Munich, Germany; Department of Computer Science, University of Chicago, Chicago, IL, USA; Computer, Electrical and Mathematical Sciences and Engineering, King Abdullah University of Science and Technology, Thuwal, Saudi Arabia; KAUST Center of Excellence for Smart Health (KCSH), King Abdullah University of Science and Technology, Thuwal, Saudi Arabia; SDAIA–KAUST Center of Excellence in Data Science and Artificial Intelligence, King Abdullah University of Science and Technology, Thuwal, Saudi Arabia; HiLIFE, Institute of Biotechnology, University of Helsinki, Helsinki, Finland; Faculty of Biological and Environmental Sciences, Organismal and Evolutionary Biology Research Programme, University of Helsinki, Helsinki, Finland; Department of Computer Science, Saint Louis University, St. Louis, MO, USA; MAP/BARseq Core Facility, Cold Spring Harbor Laboratory, Cold Spring Harbor, NY, USA; Department of Molecular Medicine, University of Padova, Padova, Italy; Department of Computer Science, Jamia Millia Islamia, New Delhi, India; Faculty of Sciences, Jamia Millia Islamia, New Delhi, India; Department of Computer Science, University College London, London, UK; Department of Biological Sciences, Purdue University, West Lafayette, IN, USA; KAUST Center of Excellence for Generative AI, King Abdullah University of Science and Technology, Thuwal, Saudi Arabia; Farming Solutions and Digital, Corteva Agriscience, Hyderabad, India; Department of Computer Science, Virginia Commonwealth University, Richmond, VA, USA; Faculty of Health and Life Sciences, University of Bristol, Bristol, UK; Graduate Institute of Biomedical Electronics and Bioinformatics, National Taiwan University, New Taipei City, Taiwan; Department of Biological Chemistry, The Hebrew University of Jerusalem, Jerusalem, Israel; School of Computer Science, Fudan University, Shanghai, China; Department of Computer Science, University of Miami, Coral Gables, FL, USA; Department of Medical Oncology, Erasmus MC Cancer Institute, Rotterdam, Netherlands; Intelligent Systems, Delft University of Technology, Delft, Netherlands; Department of Pharmacy and Biotechnology, University of Bologna, Bologna, Italy; Computational Biology for Infection Research, Helmholtz Centre for Infection Research, Brunswick, Germany; European Molecular Biology Laboratory, European Bioinformatics Institute, EMBL-EBI, Wellcome Genome Campus, Hinxton, UK; Department of Informatics, FICH-UNL, Research institute for Signals, Systems and Computational intelligence, sinc(i), CONICET/UNL, Santa Fe, Argentina; Molecular Cell Biomechanics Laboratory, Departments of Bioengineering and Mechanical Engineering, University of California Berkeley, Berkeley, CA, USA; Graduate School of Information Sciences, Tohoku University, Sendai, Japan; Genomics Molecular and Data Science, Corteva Agriscience, IN, USA; WPI-AIMEC, Tohoku University, Sendai, Japan; Department of Biology, The Hebrew University of Jerusalem, Jerusalem, Israel; Department of Computer Science, Centre for Systems and Synthetic Biology, Royal Holloway, University of London, Surrey, UK; School of Applied Mathematics, Fundação Getúlio Vargas, Rio de Janeiro, Brazil; Huawei Galois Lab, Huawei Technologies France SASU, Boulogne-Billancourt, France; ESAT-STADIUS, KU Leuven, Leuven, Belgium; Software and Information Systems Engineering, Ben-Gurion University of the Negev, Beer Sheva, Israel; Department of Computer Science, National University of Computer and Emerging Sciences, Islamabad, Pakistan; School of Computing and Data Science, Oryx University (OU) with Liverpool John Moores University, Doha, Qatar; Department of Ecology and Evolution, University of Lausanne, Lausanne, Switzerland; SIB Swiss Institute of Bioinformatics, Lausanne, Switzerland; Center for Computational Biology, Flatiron Institute, New York, NY, USA; Farming Solutions and Digital, Corteva Agriscience, IA, USA; Department of Electrical and Computer Engineering, Texas A&M University, College Station, USA; FICH, Research Institute for Signals, Systems and Computational Intelligence (sinc(i)); Universidad Nacional del Litoral (UNL), CONICET; The Rachel and Selim Benin School of Computer Science and Engineering, The Hebrew University of Jerusalem, Jerusalem, Israel; Institute of Biotechnology, University of Helsinki, Helsinki, Finland; Pink Data Analytics, Croatia; Signal Theory, Telematics and Communications, University of Granada, Granada, Spain; Interuniversity Institute of Bioinformatics in Brussels, ULB-VUB, Brussels, Belgium; Department of Computer and Information Sciences, Temple University, Philadelphia, PA, USA; School of Computing and Mathematical Sciences, Birkbeck, University of London, London, UK; Department of Biology, University of Miami, Coral Gables, FL, USA; Sylvester Comprehensive Cancer Center, University of Miami, Coral Gables, FL, USA; School of Natural Sciences, University of Kent, Canterbury, Kent, UK; Environmental Bioinformatics, SIB Swiss Institute of Bioinformatics, Lausanne, Switzerland; The Faculty of Engineering, University of Technology Tallinn, Tallinn, Estonia; University of Southern Denmark, Denmark; School of Mathematics and Statistical Sciences, National University of Ireland Galway, Galway, Ireland; Institute of Science and Technology for Brain-Inspired Intelligence and MOE Frontiers Center for Brain Science, Fudan University, Shanghai, China; School of Statistics and Data Science, Nankai University, Tianjin, China; Yunes Foundation for Research on Aging, San Francisco, CA, USA; CAS Key Laboratory of Quantitative Engineering Biology, Shenzhen Institute of Synthetic Biology, Shenzhen Institutes of Advanced Technology, Chinese Academy of Sciences, Shenzhen, China; Department of Computer Science, School of Computing, National University of Singapore, Singapore; Department of Biochemistry, Yong Loo Lin School of Medicine, National University of Singapore, Singapore; Cancer Science Institute of Singapore, National University of Singapore, Singapore; Computer and Information Sciences Department, St. Ambrose University, Davenport, IA, USA; College of Information Engineering, Zhejiang University of Technology, Zhejiang, China; College of Artificial Intelligence, Nanjing Agricultural University, Jiangsu, China; Department of Computer Science, Florida Memorial University, Miami Gardens, FL, USA; Chalmers E-commons, Chalmers University of Technology, Göteborg, Sweden; Chair for Bioinformatics, Technical University of Munich, Munich, Germany; Department of Biological Sciences, Carnegie Mellon University, Pittsburgh, PA, USA; Department of Computational Biology, Carnegie Mellon University, Pittsburgh, PA, USA; Department of Plant and Microbial Biology and Center for Computational Biology, University of California, Berkeley, CA, USA; Department of Biomedical Informatics, School of Medicine, University of Colorado Anschutz Medical Campus, Aurora, CO, USA; Center for Information Technology, National Institutes of Health, Bethesda, MD, USA

## Abstract

**Background:** The Critical Assessment of Functional Annotation (CAFA) is a community effort held to understand the field of computational protein function prediction. Every three years, since 2010, the organizers initiate an experiment to collect function predictions on a large set of proteins and then evaluate the performance of predicting methods on a subset of proteins that have accumulated experimental annotations between the submission deadline and the evaluation time. CAFA provides an independent and rigorous assessment of the current state of the art, thus leveling the playing field, highlighting successes, revealing bottlenecks, and offering a forum for the exchange of ideas in protein science. Here, we report the results of the fourth CAFA experiment (CAFA4).

**Results:** CAFA4 featured the participation of 148 methods from 70 research groups on a total of 46,205 unique proteins over a 5-year annotation accumulation phase, the longest in any CAFA. In a comparison across CAFA2-CAFA4 methods, the prediction of Gene Ontology (GO) terms has clearly improved across all three GO aspects and traditional evaluation settings. While not achieving the first rank, several CAFA2 and CAFA3 methods featured in the top ten methods in many evaluations, suggesting that earlier methods still hold relevance. The performance is weaker in the newly introduced “partial knowledge” evaluation category (proteins with experimental annotations before submission deadline that gained additional annotations in the same GO aspect during the annotation accumulation phase), highlighting the need for a new class of methods. The rankings of the methods were stable over the years in traditional evaluation settings, but less so in the new partial knowledge evaluation. Overall, the field continues to progress with some influx of new participants. Sustained efforts will be necessary to substantially advance it.

## 1 Introduction

The Critical Assessment of Functional Annotation (CAFA) [1–3] is an ongoing effort to periodically evaluate protein function prediction algorithms [4–31]. A rigorous and independent assessment is designed to establish the state of the art in the field, showcase promising methodological developments, identify bottlenecks, and track progress, while allowing su”cient time between the rounds of assessments for the development and maturation of new methods. The meetings associated with CAFA, typically the International Conference on Intelligent Systems for Molecular Biology (ISMB), additionally provide a forum for examining the state of the field by bringing together computational biologists, experimental scientists, and biocurators [32, 33].

Since its inception, CAFA has driven the development of new assessment metrics, provided benchmark datasets for evaluation, defined baseline performance, tracked progress in the field, and facilitated experimental discoveries. CAFA1, conducted in 2010-2011, reported a major finding that machine learning methods significantly outperformed sequence-similarity-based searches [34] that were routinely used by experimental scientists in their workflows [1]. CAFA2, conducted in 2013-2014, expanded the number of ontologies, benchmark proteins, and assessment scenarios [2]. It documented increased participation and a considerable improvement in performance accuracy, driven by both expanded annotation databases and methodological developments. CAFA3, conducted in 2016-2017, continued to report improved performance and also featured synergy between experiments and computation [3]. On one hand, high-throughput screens allowed unbiased performance evaluation of certain microbial functions [3], while, on the other hand, computational tools played a key role in prioritizing experimental screens to simultaneously evaluate predictions and identify novel long-term memory genes in *Drosophila melanogaster* [35].

The central idea of the CAFA challenges is to evaluate the prediction methods in a time-delayed fashion, wherein the participants are asked to make predictions on a set of proteins that lack (complete) experimental annotations at the time of algorithm development and submission. The prediction season ends with a submission deadline after which the organizers wait for the experimental annotations to accumulate and then evaluate the performance of submitting methods on newly accumulated annotations. The challenge included the prediction of Gene Ontology (GO) terms [36] from the Molecular Function (MF), Biological Process (BP) and Cellular Component (CC) aspects, the three categories of GO. In CAFA2, performance assessment additionally included Human Phenotype Ontology [37] term predictions. The evaluation strategies and metrics carried out in CAFA have been scrutinized [38–40] and have, overall, become widely accepted [41–43]. Further organizational details are described in [32].

In this paper we report on the 4^th^ round of the CAFA experiment (CAFA4). The challenge was announced in October 2019 and the predictions were collected in February 2020. Three preliminary evaluations, with benchmark datasets assembled in 2020, 2021 and 2024 were conducted and disseminated at ISMB 2020, ISMB 2022 and ISMB 2024. Compared to the earlier CAFA rounds, here we allowed for a substantially longer wait time for accumulation of experimental annotations to provide a prospective comparative assessment on the largest set of proteins to date. CAFA4 features performance assessments using an expanded set of metrics, additional focus on disordered proteins, as well as a new type of assessment that allowed us to evaluate predictions in GO aspects that previously already contained experimental annotations, a setting expected to dominate in the future. The results suggest a continued performance increase across most aspects, metrics, and evaluation settings, although we have not observed breakthrough performance. The evaluation framework and results were previously described in Ramola’s doctoral dissertation [44].

## 2 Methods

### 2.1 Challenge setting

We used a time-delayed (prospective) challenge setting as in previous CAFA rounds [1–3]. Since the evaluation is conducted only on annotations collected after the challenge submission period, there is little room for information leakage between the data available before the challenge deadline and the ground truth used for evaluation. This gives method developers additional freedom to use input data however they choose.

At time *t*_𝒯_, we released a set of target proteins 𝒯 (|𝒯| = 97,999) for the challenge. At the submission deadline time *t*_*s*_, we closed the challenge and collected all submission files. At a later time point *t*_ℬ_, we then collected a subset of proteins from 𝒯 that accumulated new experimental annotations after *t*_*s*_ in each evaluation category to create a labeled benchmark set ℬ on which the methods were evaluated. Each evaluation category may be associated with a different benchmark set ℬ, and in evaluations repeated at different time points, the subsequent datasets are not necessarily supersets of the previous benchmark sets due to some retracted annotations from UniProt [45]. The time point for the final evaluation was selected to be *t*_ℬ_ = March 7, 2025, denoted by 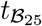 from here onwards, and the benchmark dataset at 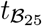 is denoted by ℬ_25_. Figure 1a illustrates the challenge timeline and four additional evaluation time points, *t*_ℬ_ = February 17, 2021, *t*_ℬ_ = September 16, 2022, *t*_ℬ_ = February 2, 2023 and *t*_ℬ_ = February 9, 2024, selected to assess how the top methods performed over time. For simplicity, these will be denoted by 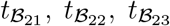, and 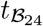,the associated benchmark datasets will be denoted by ℬ_21_, ℬ_22_, ℬ_23_ and ℬ_24_, respectively. The dataset and implementation details are provided in Supplementary Materials S1.

**Figure 1:**
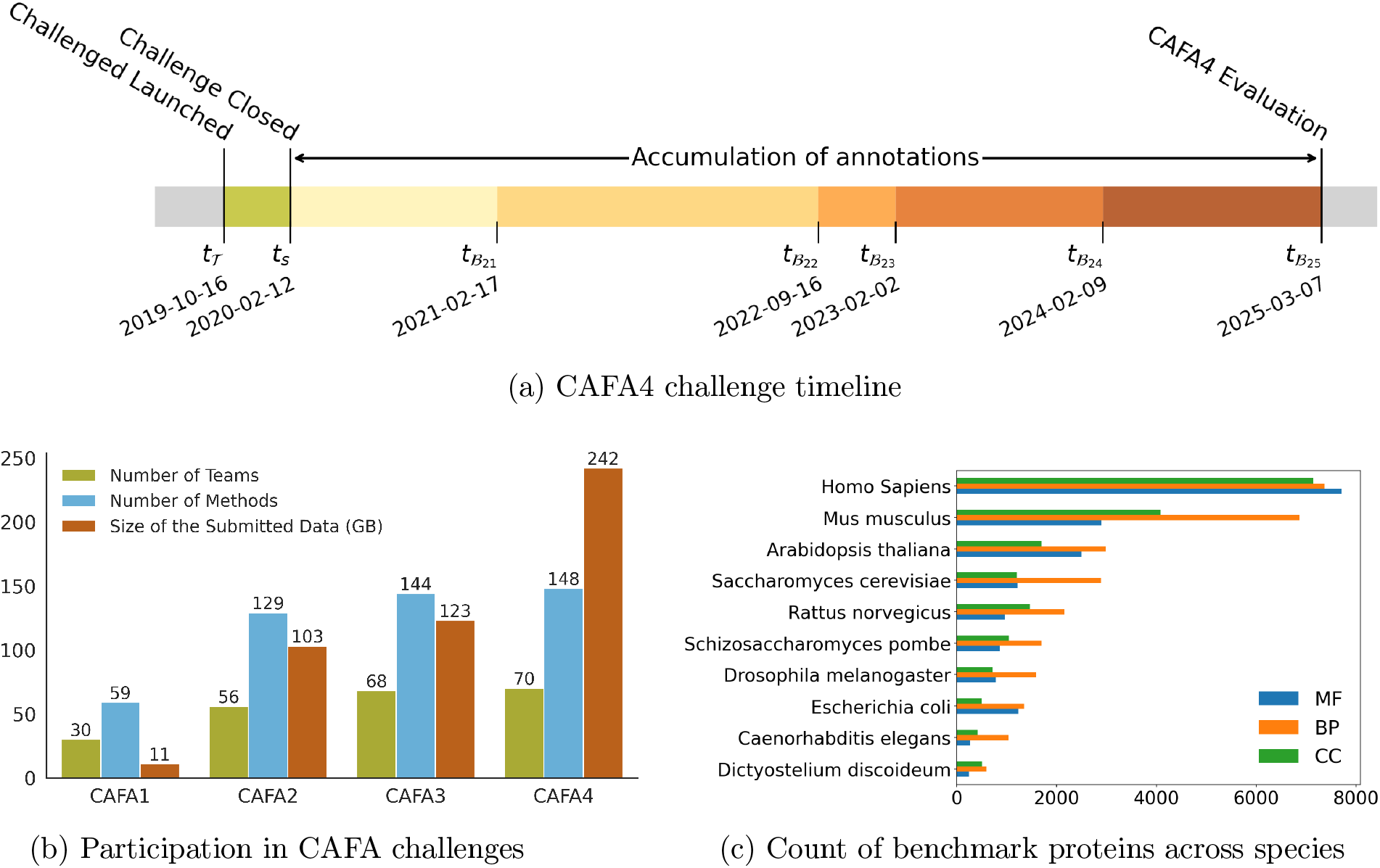
(a) Timeline of the CAFA4 challenge. The predictions could be submitted between October 16, 2019 and February 2, 2020. The evaluation was performed at five time points based on annotations collected from five UniProt releases: 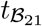 (UniProt Release 2021-02-17), 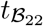 (UniProt Release 2022-09-16), 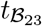 (UniProt Release 2023-02-02), 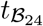 (UniProt Release 2024-02-09) and 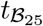 (UniProt Release 2025-03-07). (b) Participation numbers in CAFA challenges. (c) The breakdown of the proteins in the benchmark dataset across species for the three GO aspects: Molecular Function (MF), Biological Process (BP), and Cellular Component (CC).

Each row in the list of predicted annotations consisted of a protein’s identifier (ID), a predicted GO term, and a probabilistic score in the interval (0, 1], estimating the strength of the protein-term association. Each team could submit predictions from at most three different methods, from which we selected the method with the highest performance on each benchmark set to represent the team. In CAFA4, the prediction tasks involved predicting protein-term associations in all three aspects of GO [36] and one separate category involved predicting the function of disordered proteins. A total of 148 methods from 70 teams participated in CAFA4 (Figure 1b), of which 144 were evaluated. Figure 1b also shows an increase in the number of participating teams and data size over the years. The number of proteins in the ten main benchmark species and the three GO aspects are shown in Figure 1c. The evaluation of disordered proteins was performed according to the standard CAFA assessment framework; see details in Supplementary Materials S2.

### 2.2 Benchmark Datasets

We report the evaluation of the CAFA4 prediction methods on the annotations gained at 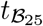 and a year-wise comparison of the top 10 methods at 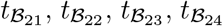 and 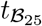. We also investigate the relative standing of the CAFA2, CAFA3, and CAFA4 methods on the 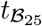 benchmark, with CAFA1 methods excluded in part due to more substantial changes of the GO graphs after 2010. For each evaluation, there are three settings, as shown in Figure 2a. The benchmark for disordered proteins was constructed using the DisProt database [46] that collected annotations between September 2019 and June 2025.

**Figure 2:**
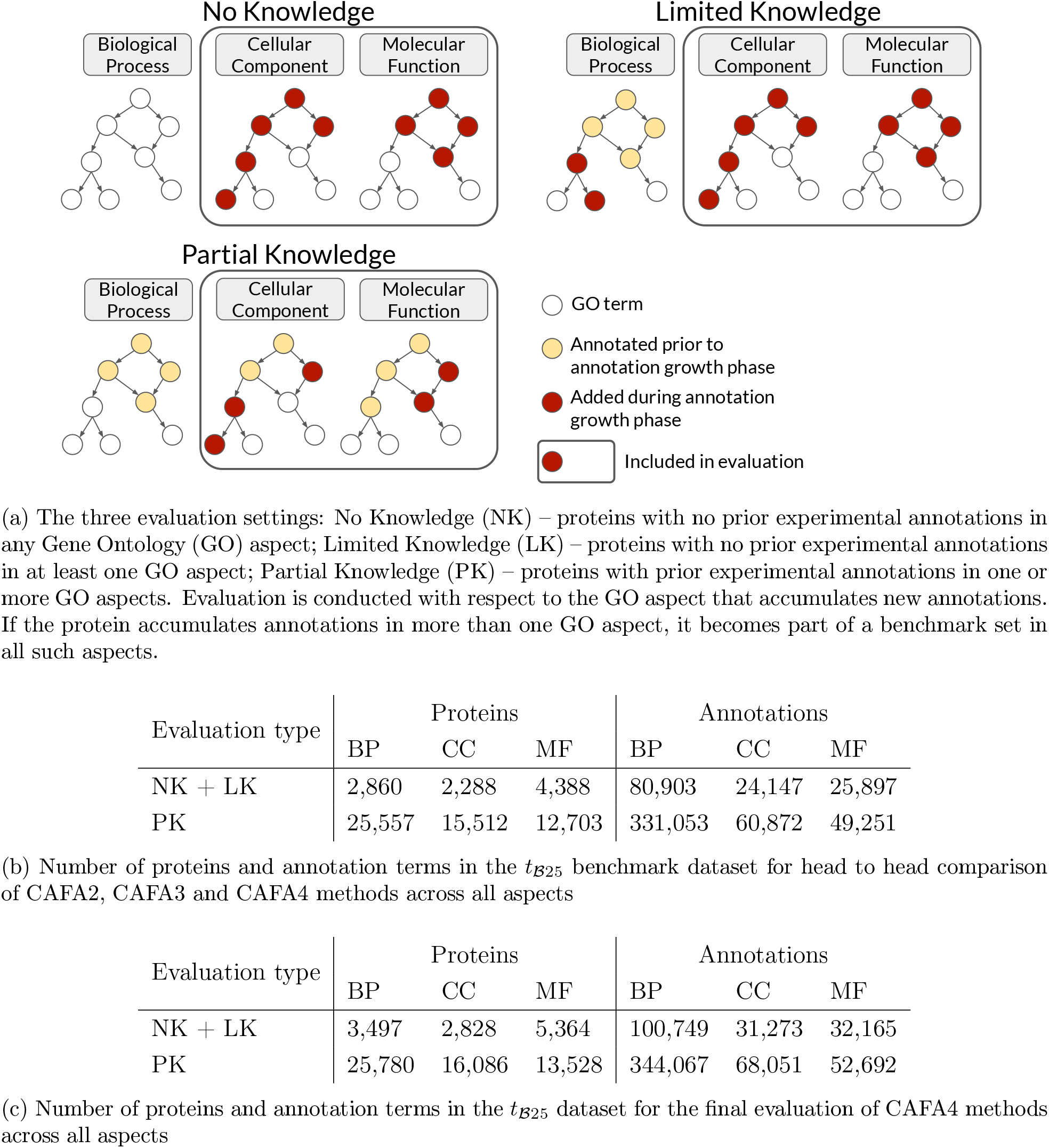
Evaluation types and benchmark datasets

#### 2.2.1 Preprocessing the annotations

The raw UniProt annotations were first filtered to only include those with experimental evidence codes (‘EXP’, ‘IDA’, ‘IPI’, ‘IMP’, ‘IGI’, ‘IEP’, ‘TAS’, ‘IC’, ‘HTP’, ‘HDA’, ‘HMP’, ‘HGI’, ‘HEP’).

Thereafter, duplicate and negative annotations were dropped and the “protein binding” term in the MF aspect (GO:0005515) was excluded. The UniProt identifier ‘Object ID’ was mapped to the respective CAFA target ID, for each benchmark protein.

#### 2.2.2 The evaluation settings

The evaluation was carried out in three settings shown in Figure 2a: (i) the No Knowledge (NK) setting includes the proteins that did not have any annotations before *t*_0_ and accumulated annotations by *t*_ℬ_ in the evaluation GO aspect, (ii) the Limited Knowledge (LK) setting includes proteins that did not have any annotations before *t*_0_ in the evaluation GO aspect, that contained at least some annotations in any of the other two GO aspects, and accumulated annotations in the evaluation GO aspect by *t*_ℬ_, and (iii) the Partial Knowledge (PK) setting includes proteins that had annotations in the evaluation GO aspect at *t*_0_, regardless of other GO aspects, and gained more annotations in that aspect by *t*_ℬ_. The PK evaluation is new in CAFA. It is expected to become a major assessment category in the future, as a large and increasing fraction of proteins, especially in model organisms, will have at least some experimental annotations. We recognize that the terminology for the three assessment scenarios is imperfect.

### 2.3 Evaluation

We used per-protein evaluation, as before [1–3]. The per-protein prediction setting refers to the task of predicting all annotations for a given target protein (structured-output classification). An ideal prediction method, say one that outputs a score of 1 for all correct terms and 0 otherwise, would achieve the maximum performance.

The main metrics for evaluating the per-protein prediction relied on computing precision-recall (pr-rc) curves and ru-mi semantic distance curves, with *F*_max_ and *S*_min_ used respectively to rank the methods [1–3]. Precision (*pr*), recall (*rc*), and *F*_max_ were estimated as follows. Consider a protein *i* from ℬ and let *P*_*i*_(τ) be the set of terms that have predicted scores greater than or equal to τ, 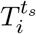 be the set of experimentally determined terms at time *t*_*s*_, and 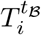 be the set of experimentally determined terms at time *t*_ℬ_, with 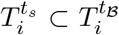. Both predicted and experimental annotations were propagated as in previous CAFAs [1–3]. In contrast to previous CAFAs, here we explicitly define macro (*M*) and micro (*µ*) versions of precision and recall, which are subsequently used to establish *F*_max_.

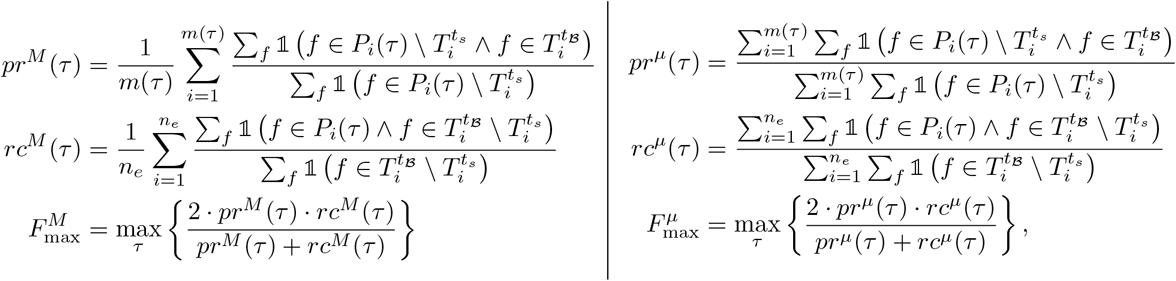

where *m*(τ) is the number of sequences with at least one predicted score greater than or equal to τ, 𝟙(*·*) is an indicator function, and *n*_*e*_ is the number of targets used in a particular mode of evaluation. In the “complete” evaluation mode *n*_*e*_ = |ℬ| = *n*, the number of benchmark proteins, whereas in the “incomplete” evaluation mode *n*_*e*_ = *m*(0); i.e., the number of proteins that were chosen to be predicted using the particular method. For each method, we refer to *m*(0)*/n* as the *coverage* (*C*) which gives the fraction of benchmark proteins on which the method made predictions. All reported evaluations in this work refer to the complete mode.

By selecting proteins for which 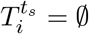, one can obtain benchmark sets that correspond to the union of what was referred to in previous rounds of CAFA as “no knowledge” and “limited knowledge” benchmarks, depending on each protein’s annotations at time *t*_*s*_ in the GO aspects other than the aspect used for evaluation. By selecting proteins for which 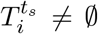, one obtains a new type of evaluation in CAFA, “partial knowledge”, again potentially split into two groups depending on its experimental annotations in the other GO aspects at time *t*_*s*_.

The remaining uncertainty (*ru*), misinformation (*mi*), and *S*_min_ only have the micro version and are defined as follows:

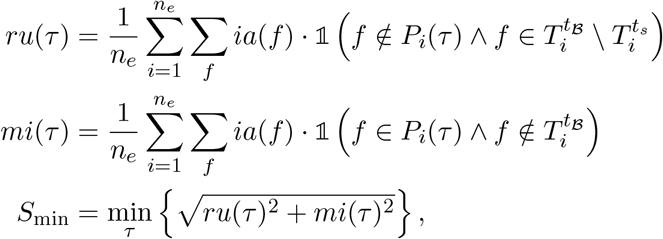

where *ia*(*f*) is the information accretion [47, 48] of the GO term *f* given UniProtKB [45] as a reference database of experimentally annotated proteins. It is estimated in a maximum likelihood manner as the negative binary logarithm of the conditional probability that the term *f* is present in a protein’s annotation given that all its parent terms are also present. Observe that here, *n*_*e*_ = *n* in the full evaluation mode and *n*_*e*_ = *m*(0) in the partial evaluation mode applies to both *ru* and *mi*.

### 2.4 Comparative evaluation over different CAFA challenges

The head-to-head evaluation of methods from all rounds of CAFA was carried out using all deposited predictions from CAFA2 [2] and CAFA3 [3] and predictions submitted during CAFA4. A separate ontology of common terms among four different versions of GO was created for this purpose. Naive [13] and BLAST [49] baseline methods from previous years were similarly provided to help us disentangle the amount of progress due to larger training data versus the progress due to methodological developments.

### 2.5 Statistical confidence

Confidence intervals from 2.5 to 97.5th percentile, in all performance evaluations, were computed using bootstrapping with 1,000 iterations [50].

## 3 Results

### 3.1 Comparative evaluation across CAFA2, CAFA3 and CAFA4

To assess improvements in protein function prediction methods across the CAFA challenges, we evaluated the methods submitted to CAFA2 [2], CAFA3 [3], and CAFA4 on a benchmark dataset of targets that were common in the three challenges. The number of common proteins and GO terms across the three GO aspects in the NK + LK (combined No Knowledge and Limited Knowledge) and Partial Knowledge (PK) categories is summarized in Figure 2b. The CAFA1 [1] methods were not included in this head-to-head comparison because of ontology changes and also because CAFA1 methods did not feature in the top 12 methods in the head-to-head comparison of CAFA1, CAFA2, and CAFA3 methods in any GO aspect in the CAFA3 [3] evaluation, showing that the field had advanced substantially since CAFA1 at the time. The comparison was done based on 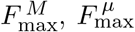 and *S*_min_ as described above. The comparisons based on 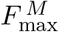, across the three GO aspects, in the NK + LK category is shown in Figure 3a and the PK category is shown in Figure 3b. The performance of all CAFA2, CAFA3 and CAFA4 methods on the common benchmarks is summarized in Supplementary Figures S3.3, S3.4, S3.5 and S3.6. The performance of the top 10 CAFA methods including the precision-recall curves are shown in Supplementary Figures S3.7, S3.8, S3.9, S3.10, S3.11, S3.12, S3.13, S3.14, S3.15, S3.16, S3.17 and S3.18.

**Figure 3:**
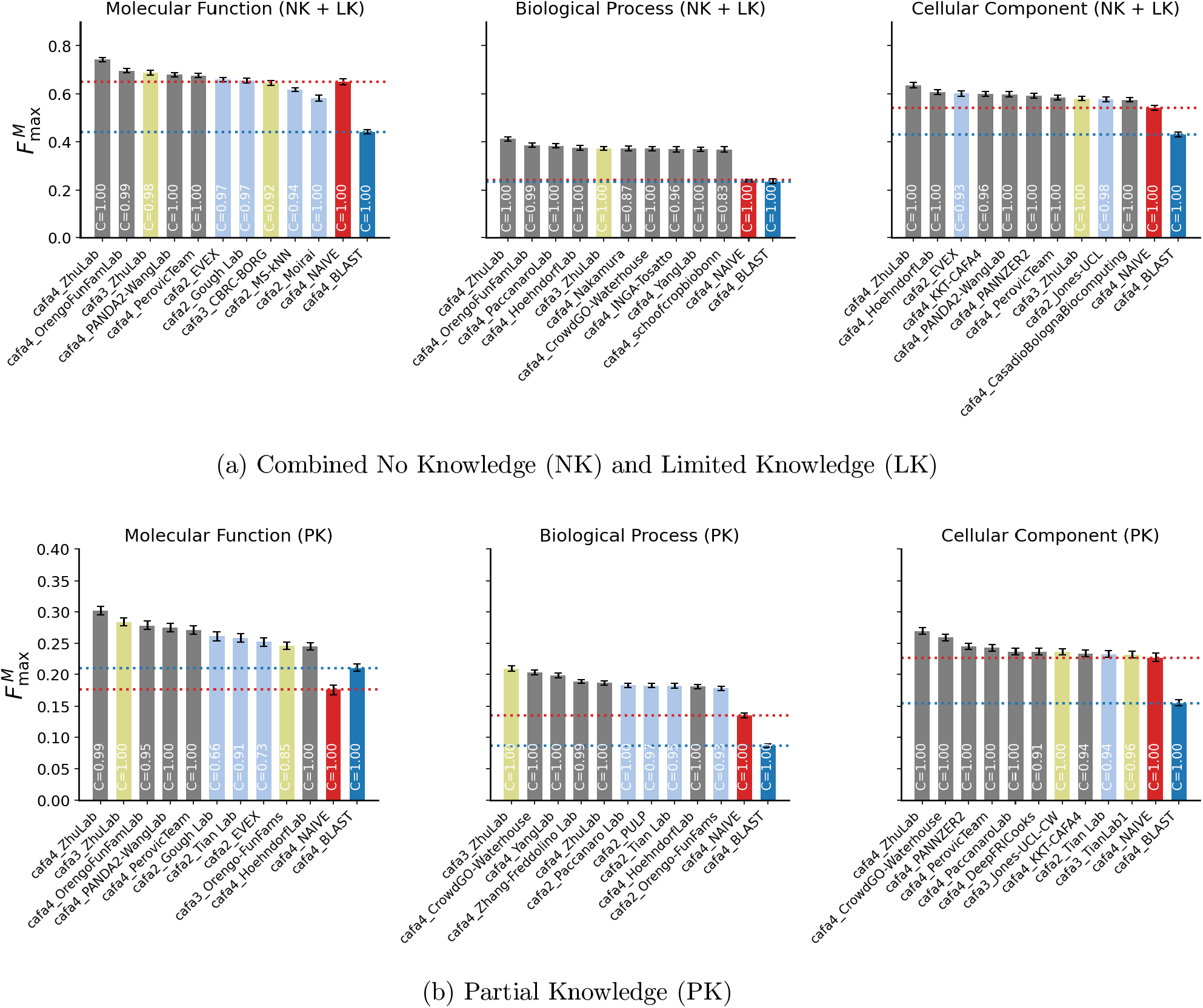
The performance of top 10 methods across CAFA2 (light blue), CAFA3 (light green) and CAFA4 (light gray) based on 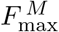 on the ℬ_25_ benchmark dataset for (a) the combined No Knowledge (NK) and Limited Knowledge (LK) dataset, and (b) the Partial Knowledge (PK) dataset. Evaluations are presented for the three Gene Ontology (GO) aspects: Molecular Function, Biological Process, and Cellular Component. Coverage (C) indicates the fraction of benchmark proteins for which each method made predictions.

The top CAFA4 methods outperformed the top CAFA2 and top CAFA3 methods across all GO aspects and all evaluation metrics in the NK + LK setting. The top method in each of these settings is from the Zhu group, though the best-performing methods within that group vary. The Orengo FunFam features second based on 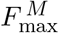 and third based on *S*_min_ in MF and BP aspects, second based on 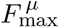 in BP, and third based on 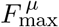 in MF. Several CAFA3 and CAFA2 methods also rank among the top 10 in every evaluation in the NK + LK setting. The top CAFA3 method featured in this evaluation is also from the Zhu lab. It is the second best performing method based on 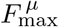 and *S*_min_ in MF and *S*_min_ in BP. The top CAFA2 method that appears in most settings is EVEX, most notably it is third based on 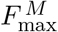 in CC. Overall, CAFA4 methods lead the top ten in every evaluation in this setting, demonstrating improvement in function prediction. However, the presence of several CAFA3 and CAFA2 methods in the top methods demonstrates their continued relevance and quality.

The PK evaluation setting is new to CAFA challenges. It has the highest number of targets in every GO aspect, and is projected to become an important evaluation category as more proteins collect annotations. As this evaluation setting had not been introduced or mentioned at the time of the CAFA4 submission deadline, the methods therefore cannot be expected to have been trained to predict deeper annotations of this setting. That notwithstanding, we assess the performance of CAFA2, CAFA3 and CAFA4 methods in this setting to understand where the field stands on the prediction of deeper annotations for proteins that were partially annotated when the challenge was announced. Compared to the NK + LK category, the scores for every metric in each aspect degrade substantially in the PK category, demonstrating the need for method development to predict annotations for this setting.

Every evaluation in this setting also showcases the Zhu lab methods, except 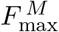 in the BP aspect, where a CAFA3 Zhu lab method performs the best. Among the CAFA4 methods, the second best-performing methods are from CrowdGO Waterhouse based on 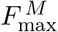 in BP and CC, the Orengo FunFam based on *S*_min_ in MF and BP, Jones-UCL based on *S*_min_ in CC, Yang lab based on 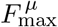 in BP and PANNZER2 based on 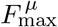 in CC. Among the CAFA3 methods, Zhu lab, Jones-UCL-CW, CBRC-BORG and Wang lab are the best-performing. Among the CAFA2 methods, Gough lab, Paccanaro lab, Tian lab, MS-kNN and Orengo FunFams are best-performing.

We also report the performance of all CAFA2, CAFA3 and CAFA4 methods on the head-to-head benchmark dataset. The distributions of 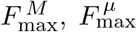 and *S*_min_ for all the methods submitted to CAFA2, CAFA3 and CAFA4 are shown in Supplementary Figures S3.3, S3.4, S3.5 and S3.6. The median score has improved in most, but not all, GO aspects for the NK + LK setting. For the PK setting, the median score for *S*_min_ improves for all aspects, and 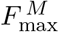 improves for the CC aspect over the course of CAFA experiments. For other settings, while the overall median has not increased in CAFA4, the top methods are from CAFA4, with the exception of 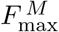 for BP in the PK setting. While the top methods in the field have improved since CAFA3 and CAFA2, overall, the field has not necessarily improved in performance in several evaluations.

### 3.2 CAFA4 Final Evaluation

To evaluate the performance of the CAFA4 methods on the benchmark dataset collected in the period since submission, they were assessed using the ℬ_25_ dataset. The number of proteins and GO terms across the three aspects in the NK + LK and PK datasets is summarized in Figure 2c. The comparisons based on 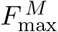, across the three GO aspects, based on the combined NK + LK dataset is shown in Figure 4a and the PK dataset is shown in Figure 4b. The performance of the top ten CAFA4 methods including the precision-recall curves are shown in Supplementary Figures S3.19, S3.20, S3.21, S3.22, S3.23, S3.24, S3.25, S3.26, S3.27, S3.28, S3.29 and S3.30.

**Figure 4:**
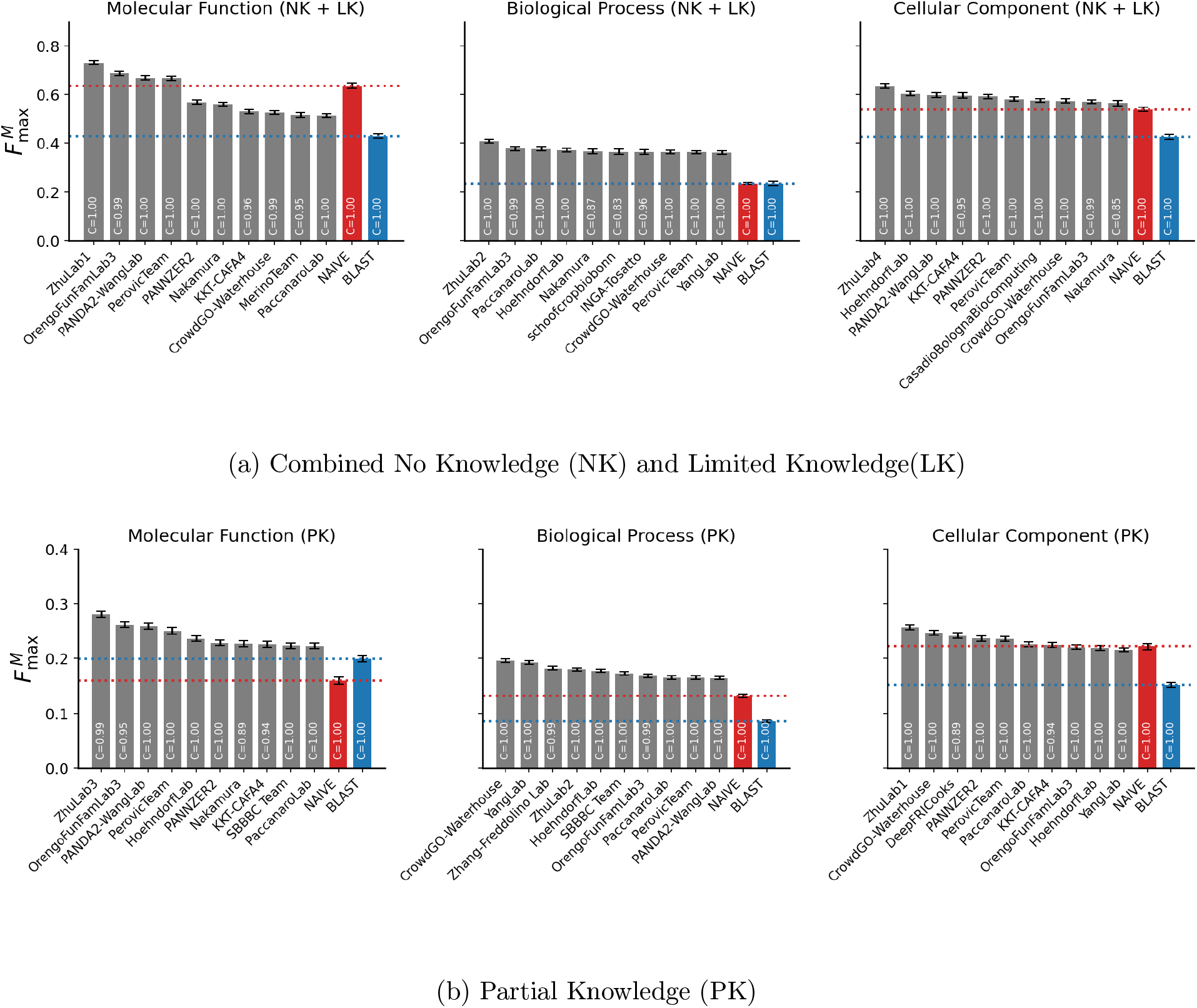
The performance of top 10 CAFA4 methods based on 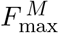 on the ℬ_25_ benchmark dataset for (a) the combined No Knowledge (NK) and Limited Knowledge (LK) dataset, and (b) the Partial Knowledge (PK) dataset. Evaluations are presented for the three Gene Ontology (GO) aspects: Molecular Function, Biological Process, and Cellular Component. Coverage (C) indicates the fraction of benchmark proteins for which each method made predictions.

For the NK + LK setting, one of the methods from the Zhu group is the best-performing across all metrics and aspects. The second best performing methods are from the Orengo FunFam group across all metrics for MF and BP aspects. In the CC aspect, the second bestperforming method is Hoehndorf lab based on 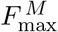, PANDA2-Wang lab based on 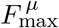 and KKT-CAFA4 based on *S*_min_. The top ten methods across GO aspects and metrics sometimes did not perform better than the baselines.

In the PK setting, one of the Zhu group methods performs the best across all but two metric-aspect combinations. The best performing method in BP is CrowdGO Waterhouse based on 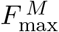, and DeepFRI Cooks in CC based on *S*_*min*_. The top ten methods outperform Naive and BLAST baselines based on 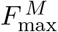 in all GO aspects. Overall, the performance of methods in the NK + LK setting is substantially better than that in the PK setting. Among those settings, the performance in MF and CC aspects is considerably better than in the BP setting.

### 3.3 Performance over the years

The CAFA4 challenge had a 5-year annotations collection phase. The evaluation dataset has evolved over the years to include more proteins and terms as shown in Supplementary Figure S3.31. To assess the stability and utility of the prediction methods over the years, we also evaluated their performance on these annual benchmark datasets from 2021-2025. The 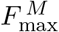 comparisons across the three GO aspects, based on combined NK + LK datasets are shown in Figure 5a and the PK datasets are shown in Figure 5b. The distributions of the performance of CAFA4 methods over the yearwise benchmarks are shown in Supplementary Figures S3.32, S3.33 and S3.34. The performance of the top ten CAFA4 methods for all the metrics and benchmarks are shown in Supplementary Figures S3.35, S3.36, and S3.37.

**Figure 5:**
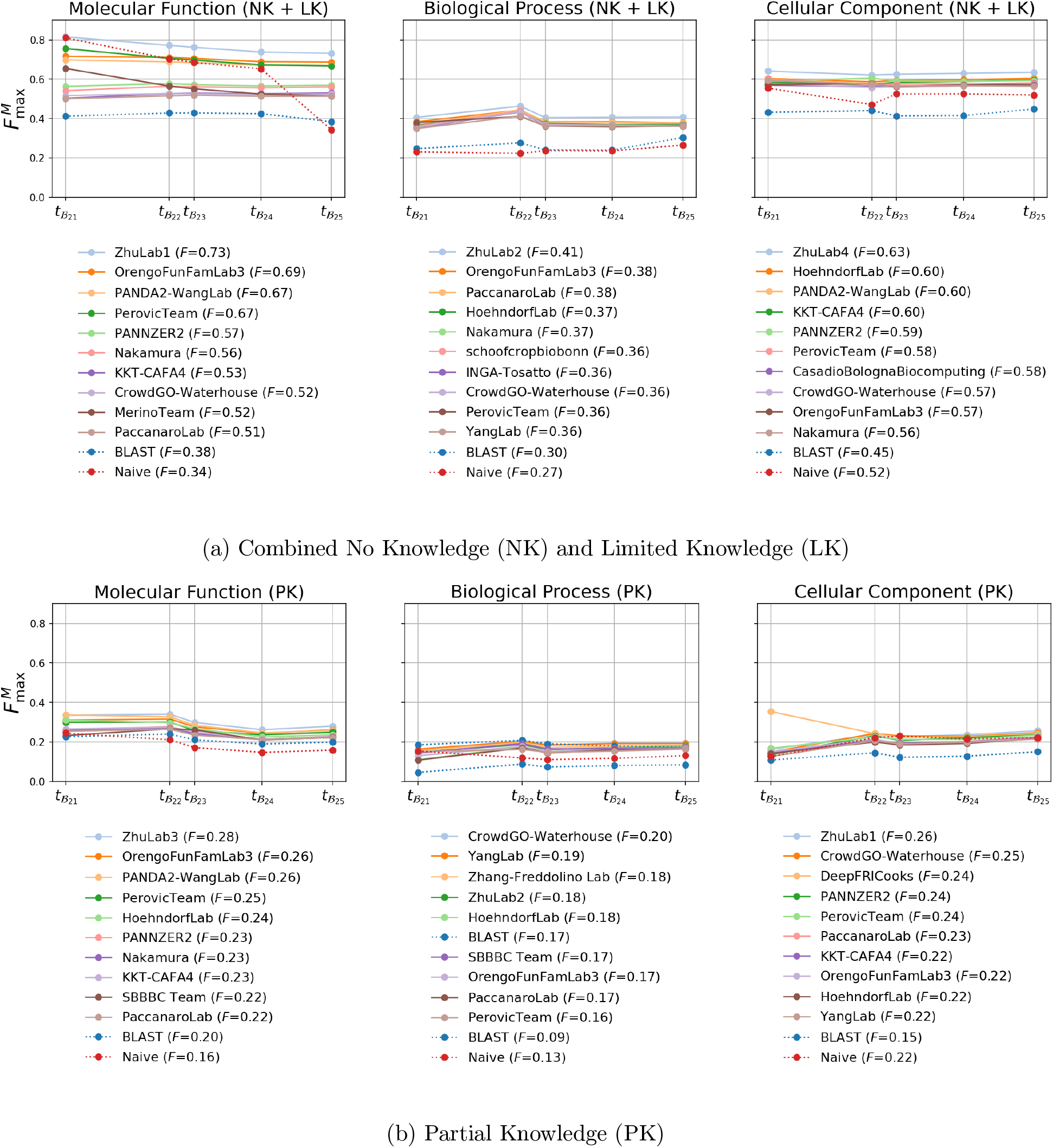
The performance of top 10 methods based on 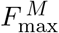 over the five yearly benchmark datasets - ℬ_21_, ℬ_22_, ℬ_23_, ℬ_24_ and ℬ_25_. The 10 best methods that feature in each category here were selected on the basis of their 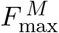 score on the ℬ_25_ benchmark. The reported *F* is the 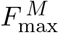 from evaluation on the ℬ_25_ benchmark dataset for (a) the combined No Knowledge (NK) and Limited Knowledge (LK) dataset, and (b) the Partial Knowledge (PK) dataset. Evaluations are presented for the three Gene Ontology (GO) aspects: Molecular Function, Biological Process, and Cellular Component. Coverage (C) indicates the fraction of benchmark proteins for which each method made predictions.

For the NK + LK setting, 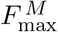 and 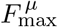 of the methods have been stable in BP and CC aspects. However, they decrease noticeably in MF. The 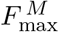 of the method from the Zhu lab drops from 0.814 at 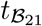 to 0.731 at 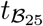 dataset. While some of the other top methods in these evaluations are relatively stable, others follow this trend of decreased performance. *S*_min_ is stable in the CC aspect, but increases slightly in the BP and MF aspects. The *S*_min_ of the Zhu lab group increases from 19.98 at 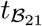 to 21.39 at 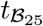 in BP and 1.61 at 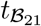 to 2.69 at 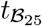 in MF. Overall, the relative ranking of the top methods, especially the first rank, remains stable over the years.

The relative rankings vary more in the PK evaluation. In the CC aspect, based on 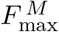 and 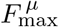, the best performing method at 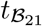 is DeepFri Cooks 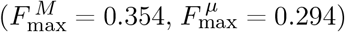. However, these scores drop substationally over the years (at 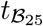, 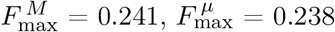). The Zhu lab score and ranking improve over the years in this evaluation, and they overtake DeepFri Cooks at 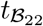. Overall, slight increases in 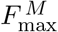 and 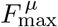 are observed for most methods in the BP and CC aspects, whereas MF is relatively stable over the years. However, *S*_min_ deteriorates for BP and CC while staying mostly stable in MF.

## 4 Discussion

Independent assessment of protein function prediction is critical for understanding the field and important for spurring further innovation [33, 43, 51]. Towards this goal, we performed a comprehensive evaluation of the CAFA2, CAFA3 and CAFA4 methods on a large benchmark dataset. We expanded the evaluation framework by including the micro metrics and introducing the partial knowledge (PK) evaluation setting. The ground truth dataset was curated from UniProt databases over a 5-year accumulation phase, the longest in any CAFA experiment. We also performed an annual evaluation of the participating methods on the growing evaluation datasets from 2021 to 2025 to assess their temporal stability and relevance.

We observed that the previous methods were outperformed by top CAFA4 methods in most evaluation settings, suggesting progress. The improvement in MF and CC aspects is more evident compared to the BP aspect, which remains a di”cult problem. In the annual evaluation from 2021-2025, the rankings of the methods were stable over the years in the NK, LK and NK+LK evaluation settings, but less so in the new PK evaluation setting. As the PK category was not introduced at the time of launch of the challenge, it is reasonable that the methods do not perform as well as the standard NK and LK settings of CAFA. It is noteworthy that the PK group has the highest number of proteins and annotations. As more proteins accumulate annotations, more annotations will fall in this group compared to NK and LK. This underscores the PK prediction problem as a new challenge for future methods.

While this work provides a comprehensive evaluation of the protein function prediction methods from the previous CAFA challenges over a long annotation accumulation period, it has some limitations. First, since the evaluation dataset is curated from UniProt databases, the biases in those databases such as more annotations from well-studied proteins and functions or potential changes in curation patterns over the years, are hard to address (some of these problems are exemplified by a relatively high accuracy of the Naive baseline method). Second, while the participants are asked to submit keywords associated with their methods, the evaluation is mostly blind to the exact machine learning methodologies or datasets used in the training. Third, the evaluation is limited to the methods that participate in the CAFA challenges, so potential strong methods that do not participate cannot be included in our assessments. Finally, the inclusion of text mining methods can be a source of data leakage. Such methods can include training on experimental annotations from publications published before the submission deadline, that were not in the corresponding UniProt Gene Ontology databases. The percentage of such annotations is estimated to be over 98%. The methods that rely on text mining can have significant initial advantage over other methods in this type of evaluation. However, as CAFA methods continue to be evaluated in subsequent CAFAs, the advantage of text mining methods disappears.

In conclusion, this work is a comprehensive and rigorous assessment of the methods that participated in CAFA2, CAFA3 and CAFA4 challenges. CAFA challenges continue to lay the groundwork for systematic evaluation and development of protein function prediction methods. While the field has evidently moved forward, the problem of function prediction remains unsolved. The method development and evaluation strategies in the field are expected to continue to grow.

