## Supplementary Information for "On the state of protein function prediction: a report on the fourth CAFA challenge"

### S1 Supplementary Information - Implementation Details

#### S1.1 Evaluation

We adapt the CAFA-evaluator [51] code base to account for PK evaluation and share the code here: https://github.com/claradepaolis/CAFA-evaluator-PK.

To evaluate predictions, we run the code as follows

```
cafaeval [onotology file] [prediction directory] [gt terms] —ia [ia file] —toi
[terms of interst] —prop fill —th_step 0.001 —no_orphans
```

For partial-knowledge evaluation,

```
cafaeval [ontology file] [prediction directory] [gt terms] —ia [ia file] —known
[terms to exclude] —toi [terms of interst] —prop fill —th_step 0.001 —no_orphans
```

#### S1.2 Data Versions and Sources

For ground truth annotations, we use annotations with GO evidence codes as listed in Table 1. Annotations with these evidence codes are also used to construct the known annotations (known at $t_s$) to be excluded when evaluating the partial knowledge setting.

Details of the gene ontology annotations (GOA) and gene ontology (GO) datasets to create each benchmark dataset are listed in Table 2.

| Code | Definition |
|---|---|
| EXP | Inferred from Experiment |
| IDA | Inferred from Direct Assay |
| IPI | Inferred from Physical Interaction |
| IMP | Inferred from Mutant Phenotype |
| IGI | Inferred from Genetic Interaction |
| IEP | Inferred from Expression Pattern |
| HTP | Inferred from High Throughput Experiment |
| HDA | Inferred from High Throughput Direct Assay |
| HMP | Inferred from High Throughput Mutant Phenotype |
| HGI | Inferred from High Throughput Genetic Interaction |
| HEP | Inferred from High Throughput Expression Pattern |
| TAS | Traceable Author Statement |
| IC | Inferred by Curator |

Table 1: Evidence codes used in CAFA 4

Table 2: Benchmark datasets and annotation resources used for propagation

| **Benchmark** | **Start Time ($t_s$)** | **Benchmark Time ($t_{\mathcal{B}}$)/GOA** | **GO** |
|---|---|---|---|
| $t_{\mathcal{B}_{21}}$ | February 12, 2020 | 2021-02-17 | 2021-02-01 |
| $t_{\mathcal{B}_{22}}$ | February 12, 2020 | 2022-09-16 | 2022-07-01 |
| $t_{\mathcal{B}_{23}}$ | February 12, 2020 | 2023-02-02 | 2023-01-01 |
| $t_{\mathcal{B}_{24}}$ | February 12, 2020 | 2024-02-09 | 2024-01-17 |
| $t_{\mathcal{B}_{25}}$ | February 12, 2020 | 2025-03-07 | 2025-02-06 |

### S2 Supplementary Information - Intrinsically Disordered Proteins

Intrinsically disordered proteins (IDPs) and intrinsically disordered regions (IDRs) lack a stable three-dimensional structure under physiological conditions. Rather than adopting a single well-defined fold, they exist as dynamic ensembles of conformations. IDPs and IDRs are widespread across eukaryotes and viruses, and it is estimated that approximately one third of all residues in the human proteome are intrinsically disordered [52].

Despite their structural plasticity, IDRs play central roles in cellular regulation. Their conformational flexibility enables transient and reversible interactions, allowing them to bind multiple partners and to adapt their shape depending on the interaction context. As a result, IDRs are key components of signaling and regulatory networks. In addition, IDRs often function as flexible linkers that maintain domains in close proximity while modulating their relative orientation and activity. Their extended and accessible conformations also make them preferred targets for post-translational modifications (PTMs), such as phosphorylation or ubiquitination, which further fine-tune protein function and enable rapid regulatory responses [53].

Given their prevalence and functional importance, accurate evaluation of IDP and IDR functions is essential for understanding cellular regulation, signaling specificity, and disease mechanisms. Many proteins involved in cancer, neurodegeneration, and viral infection rely on disordered regions for their activity, making IDPs highly relevant for both basic biology and therapeutic research. However, evaluating IDP function poses challenges that are fundamentally different from those associated with structured proteins, as function is often encoded in short, context-dependent regions rather than in a single stable fold [54].

Our current knowledge of IDP functions remains limited, largely due to the experimental challenges associated with their characterization. The transient, dynamic, and context-dependent nature of IDP-mediated interactions complicates the use of classical structural biology techniques such as X-ray crystallography or cryo-electron microscopy. Furthermore, IDP functions often depend on specific cellular conditions, binding partners, or post-translational modifications, making them difficult to isolate and study in vitro. As a consequence, functional evidence for IDPs is typically fragmented, low-throughput, and distributed across diverse experimental setups. This difficulty is reflected at the database level by the scarcity of curated functional annotations for intrinsically disordered regions [55].

One of the main resources for IDP functional annotation is the DisProt database [56], which provides manually curated information on intrinsically disordered proteins and regions supported by experimental evidence. In contrast to classical Gene Ontology (GO) annotations, which are generally assigned to entire protein sequences, DisProt associates functions with specific disordered regions, enabling a more fine-grained representation of IDP functionality. Currently, functional annotations in DisProt are derived from both the Intrinsically Disordered Proteins Ontology (IDPO) and the Gene Ontology. IDPO captures functions that are specific to intrinsically disordered regions and are not explicitly represented in GO, such as "flexible linker" (IDPO:0000033) or "self regulatory activity" (IDPO:0000057). In parallel, GO terms are used to describe molecular functions such as "protein binding" (GO:0005515) or "molecular function regulator" (GO:0098772), as well as biological processes including "localization" (GO:0051179).

#### S2.1 Methods

The evaluation of Disorder Ontology (DO) term prediction follows the standard CAFA Gene Ontology (GO) assessment framework. Although DO terms in DisProt are originally annotated at the level of specific protein regions, regional information is ignored in this challenge. All DO annotations are therefore treated as if they were associated with the full-length protein, enabling a direct comparison with CAFA-style whole-protein function prediction methods.

##### S2.1.1 Benchmarking Dataset

The benchmarking dataset was constructed using DisProt proteins that became available after the CAFA4 announcement. Specifically, proteins annotated between September 2019 and June 2025 were selected, ensuring that no annotation information for these proteins was accessible to predictors at submission time. This benchmark corresponds to the CAFA "no-knowledge" setting, in which predictors are evaluated on proteins that had no DO annotations available at the time of the challenge announcement.

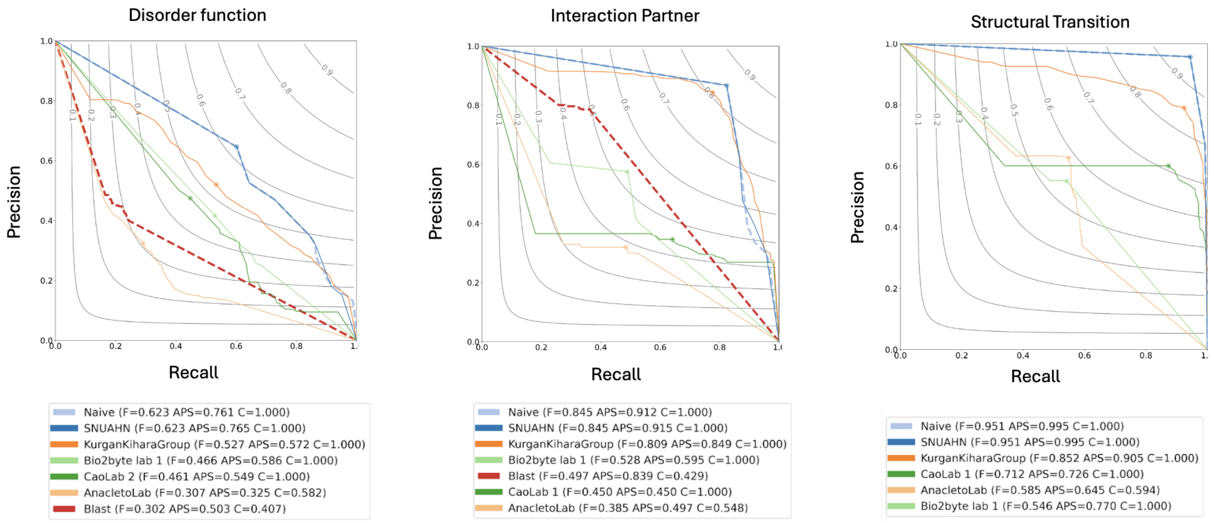


(a) Disorder function (b) Interaction partner (c) Structural transition

Figure S2.1: Precision-recall curves for intrinsic disorder challenge. Evaluation was carried out on "No knowledge" benchmark, in the full mode. $n = 354$ Disorder function; $n = 117$ Interaction partner; $n = 128$ Structural transition. The perfect prediction should have $F_{\max} = 1$, at the top right corner of the plot. The dot on the curve indicates where the maximum $F$ score is achieved.

At the time CAFA4 was announced, the Intrinsically Disordered Proteins Ontology (IDPO) was referred to as the Disorder Ontology (DO). Since then, several IDPO terms have been migrated into the Gene Ontology and removed from IDPO. To ensure a consistent evaluation framework, all IDPO and GO terms associated with newly annotated DisProt proteins were mapped back to the original DO ontology version available in September 2019.

The original DO ontology is organized into four aspects: Disorder function, Structural transition, Binding partner, Structural state. The structural state aspect was excluded from the evaluation because more than 97% of the proteins are annotated with the same term ("Disorder"), making this aspect non-informative for benchmarking purposes. Although the binding partner aspect has since been removed from IDPO because all related terms are now represented in GO, it was included in this evaluation. All binding partner annotations were retained by exploiting the one-to-one mapping between DO binding partner terms and their corresponding GO terms.

The final benchmark dataset includes: 354 proteins annotated with Disorder function terms, 177 proteins with Binding partner (interaction partner), 128 proteins with Structural transition.

##### S2.1.2 Baselines

Two baseline predictors, Naive and Blast, were constructed using only annotation data available prior to the CAFA submission deadline (January 2020), specifically from DisProt release 2019_09.

The Naive baseline assigns DO terms based on their relative frequencies in the training dataset, independently of the query protein sequence.

The Blast baseline transfers annotations from previously annotated proteins to query proteins based on sequence similarity. Query proteins were aligned against the DisProt 2019_09 dataset using BLAST [34] with default parameters. Only hits with an E-value $\leq 0.01$ were retained. Transferred annotations were weighted by the pairwise sequence identity between the query and the matched protein.

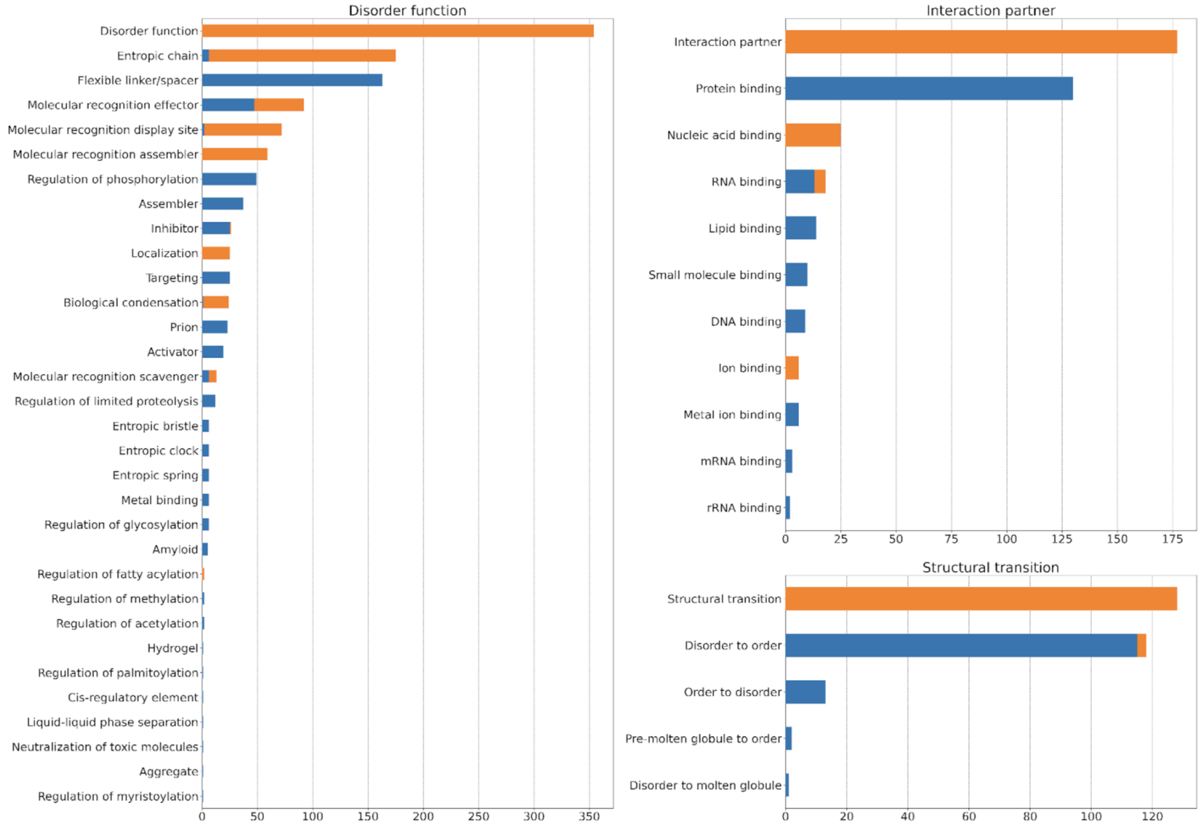


Figure S2.2: Distribution of terms in the ground truth before and after ancestors expansion. Terms are divided according to the three aspects available in the DO ontology in January 2019. Protein annotations are obtained from the DisProt database release 2025_06 after mapping IDPO and GO terms to the old DO ontology. The blue bars indicate the number of proteins directly annotated with a term, the orange bars indicate the number of proteins annotated with a term as a consequence of the ancestor expansion following the "is_a" relationship across terms.

#### S2.2 Results

All evaluations have been performed in full-mode, i.e. measuring the recall over all benchmark targets and the precision on the predicted targets at a given score threshold. The benchmark type corresponds to the CAFA "no knowledge", i.e. the evaluated proteins did not have any DisProt annotation before the submission of the predictions. Only methods that predicted at least 30% of the benchmark targets are considered. Figure S2.1 shows the performance of the top methods along with the Naive and Blast baselines. A number of methods outperform the Blast baseline while Naive is the top method for all three aspects and the precision recall curve is almost identical to the top non-baseline method. The exceptional performance of Naive can be explained by the high frequency of a limited number of terms, namely "Flexible linker/spacer" (DO:00002/IDPO:0000033), "Protein binding" (DO:00063/GO:0005515) and "Disorder to order" (DO:00050/IDPO:0000011) are associated to 163 (46.0%), 130 (73.4%) and 115 (89.8%) proteins for the "Disorder function", "Molecular interaction" and "Structural transition" aspects, respectively. The abundance of every term after ancestor expansion is provided in Figure S2.2.

The poor performance of Blast baseline, compared with the usually good performance that has been observed in classical CAFA challenges over the years, can be explained by the fact that the benchmarking dataset includes proteins that are not homologous to the proteins available in the "old" DisProt for transferring functions.

#### S2.3 Data Availability

The ground truth datasets, mapping of the predictions and baselines have been generated using Python Notebooks available at URL: https://github.com/BioComputingUP/CAFA4-IDPO.

The IDPO evaluation has been calculated using a Python implementation of the CAFA evaluation [51].

#### S3 Supplementary Information - Additional Figures

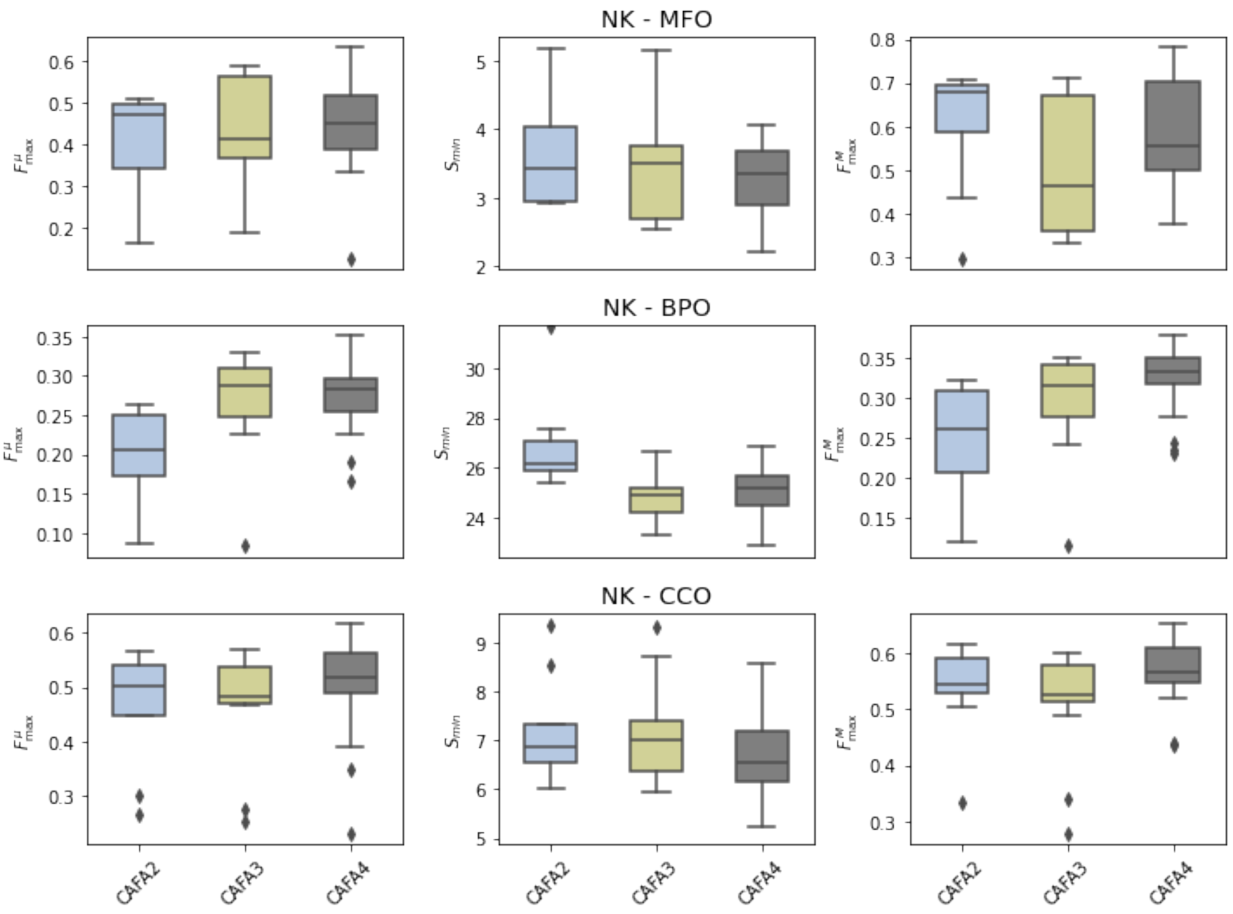


Figure S3.3: The state of the field over subsequent CAFAs. The distributions of $F^{\mu}_{\text{max}}$ on the *no knowledge* (NK) evaluation for all methods of CAFA2, CAFA3 and CAFA4 for Molecular Function ontology (MFO), Biological Process ontology (BPO), Cellular Component Ontology (CCO)

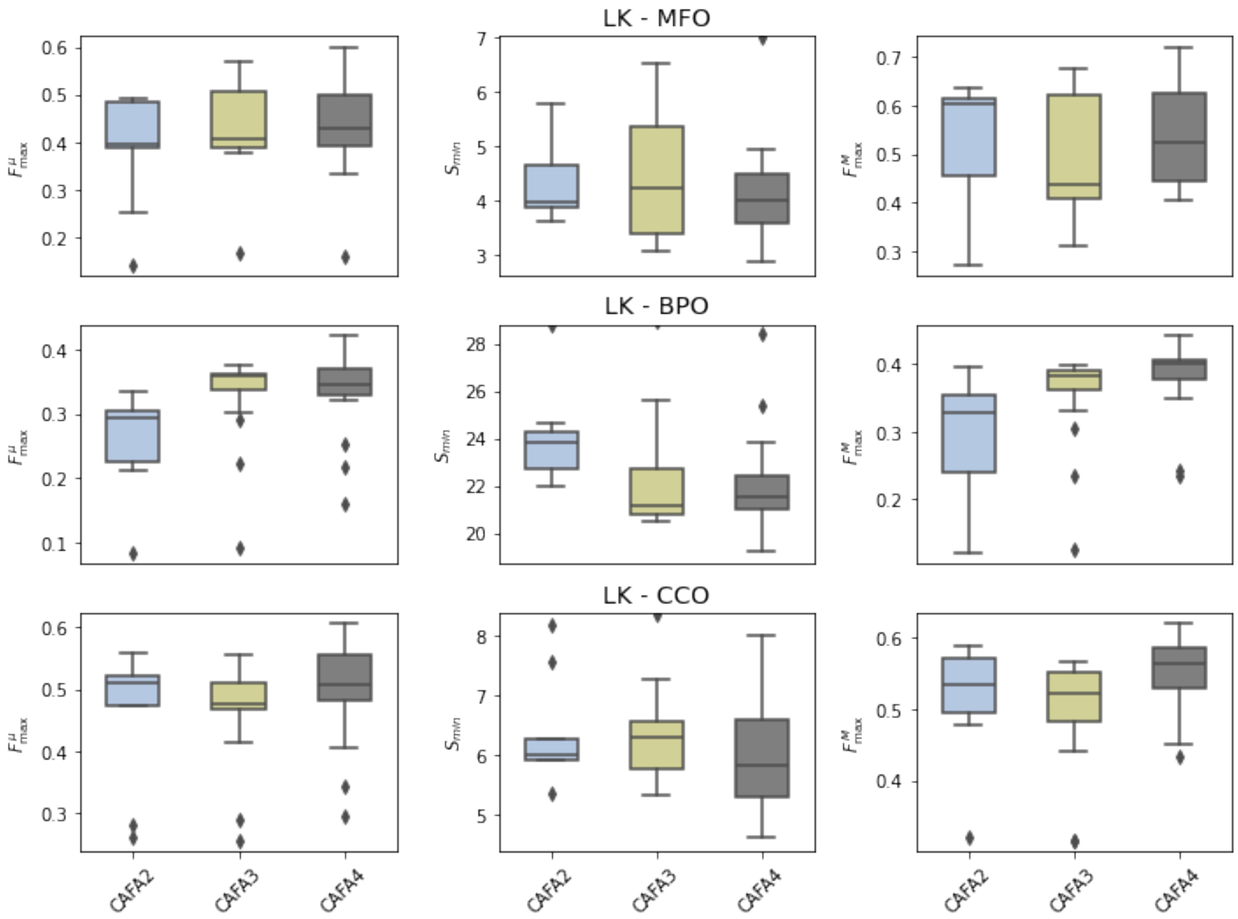


Figure S3.4: The state of the field over subsequent CAFAs. The distributions of $F^{\mu}_{\max}$ on the *limited knowledge* (LK) evaluation for all methods of CAFA2, CAFA3 and CAFA4 for Molecular Function ontology (MFO), Biological Process ontology (BPO), Cellular Component Ontology (CCO)

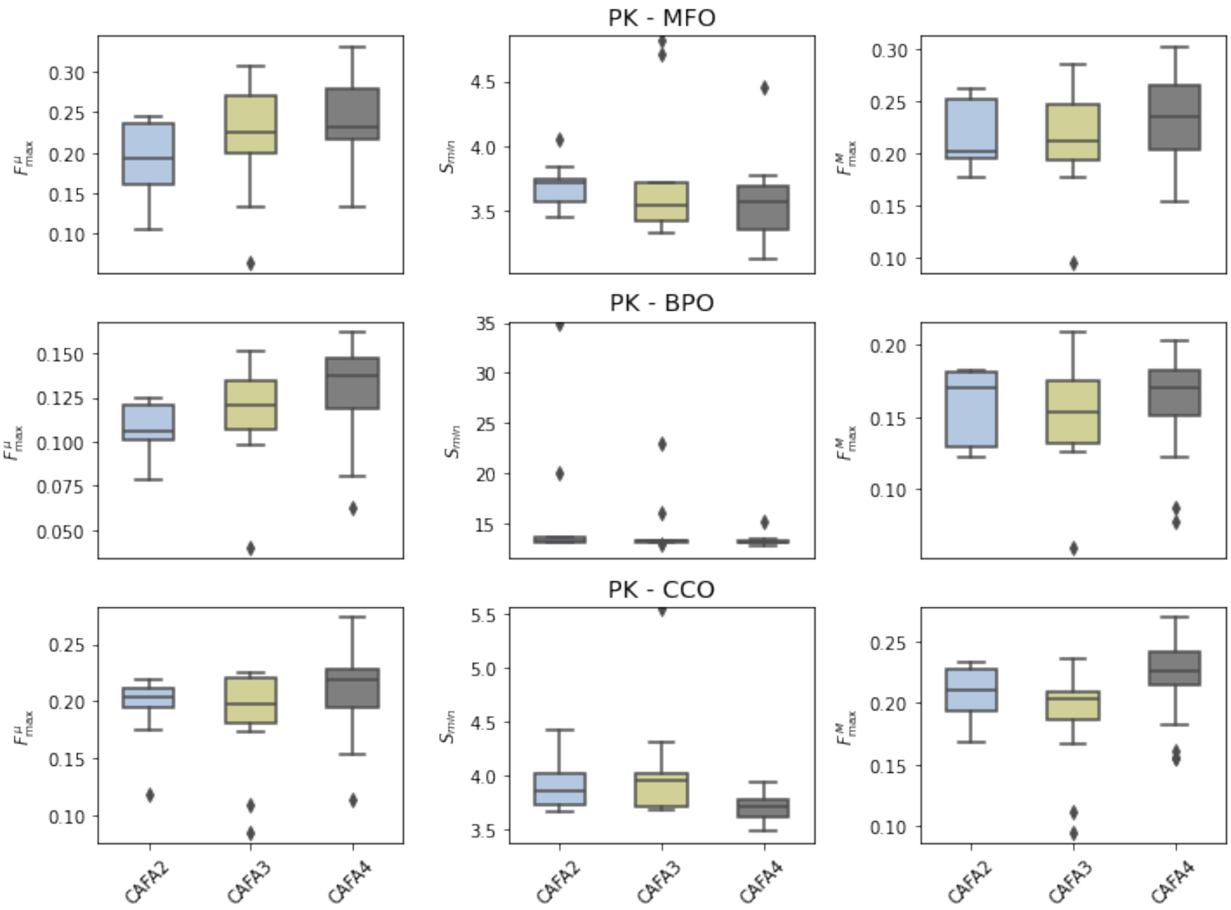


Figure S3.5: The state of the field over subsequent CAFAs. The distributions of $F^{\mu}_{\max}$ on the *partial knowledge* (PK) evaluation for all methods of CAFA2, CAFA3 and CAFA4 for Molecular Function ontology (MFO), Biological Process ontology (BPO), Cellular Component Ontology (CCO)

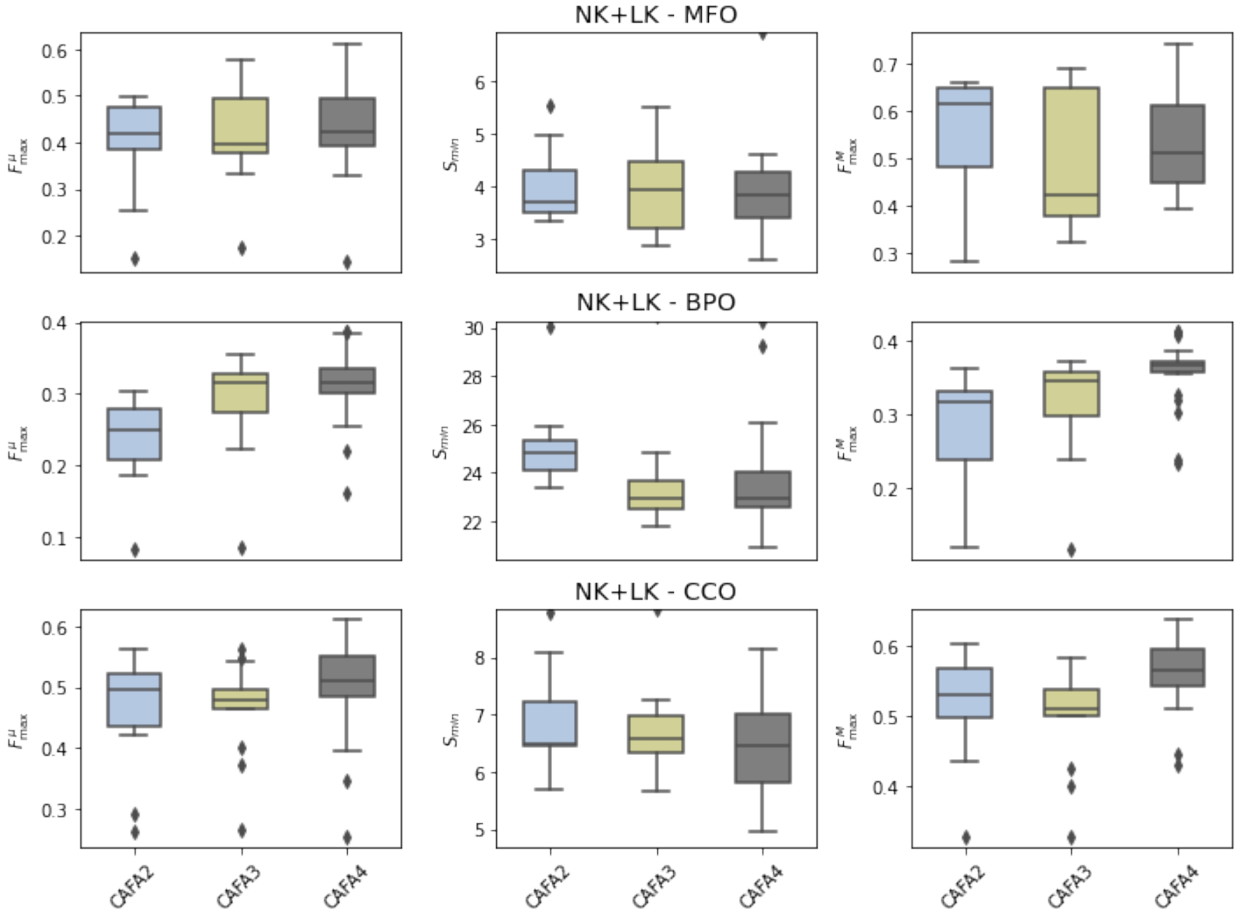


Figure S3.6: The state of the field over subsequent CAFAs. The distributions of $F^{\mu}_{\max}$ on the *no knowledge* (NK) and *limited knowledge* (LK) evaluations for all methods of CAFA2, CAFA3 and CAFA4 for Molecular Function ontology (MFO), Biological Process ontology (BPO), Cellular Component Ontology (CCO)

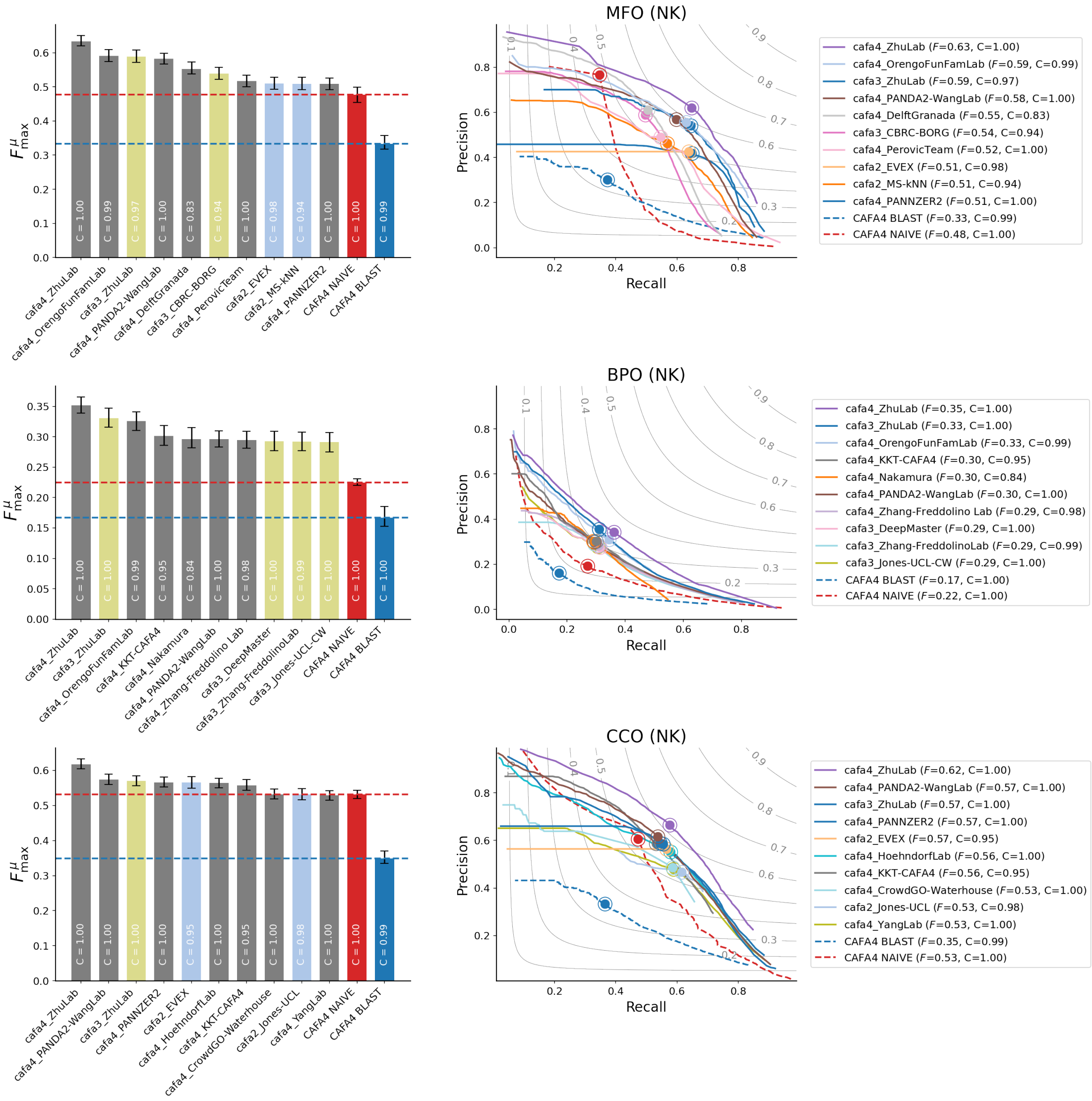


Figure S3.7: $F^{\mu}_{\max}$ scores and precision-recall curves for the head-to-head comparison of CAFA2, CAFA3 and CAFA4 methods on the *no knowledge* (NK) benchmarks

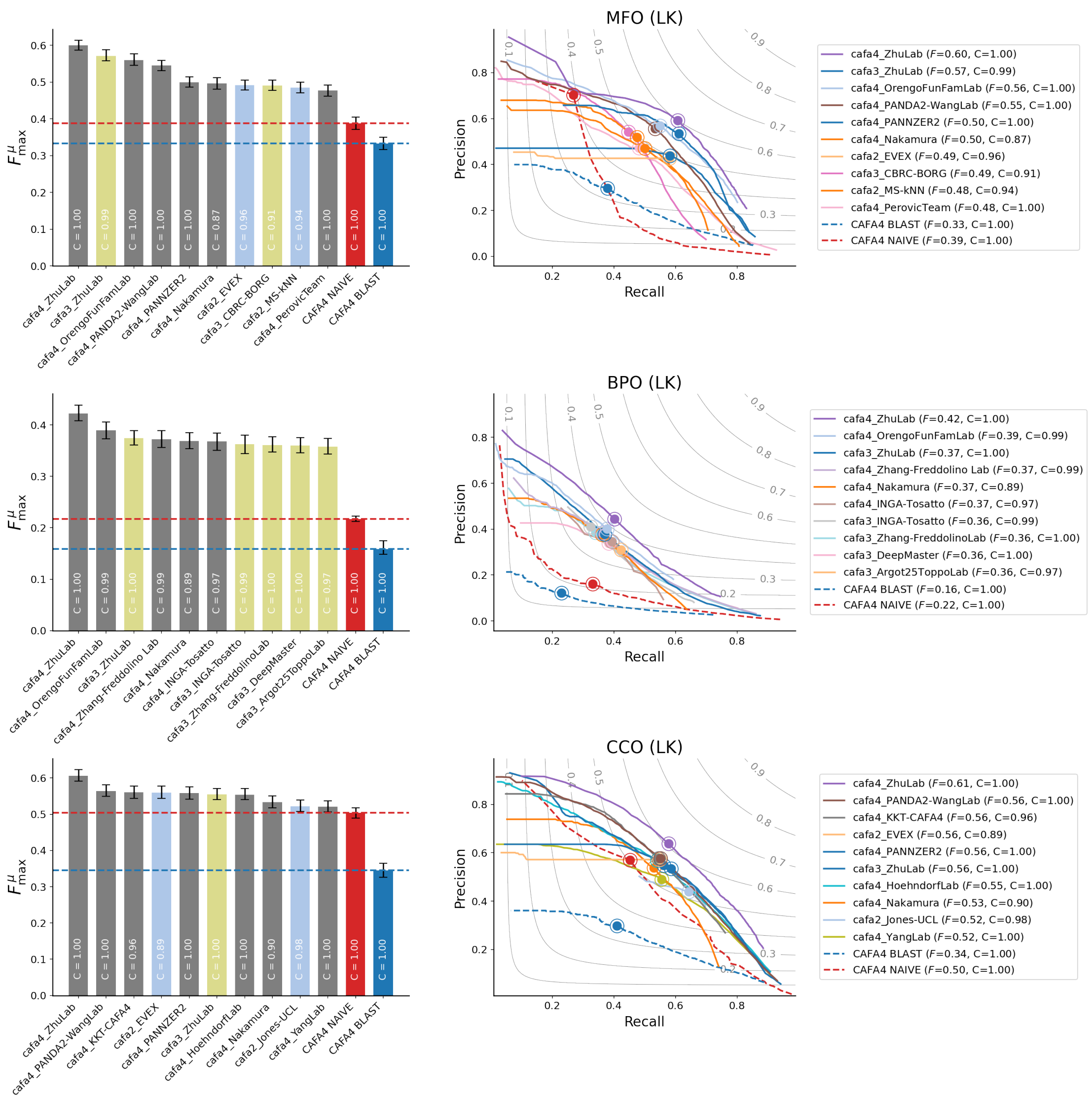


Figure S3.8: $F^{\mu}_{\text{max}}$ scores and precision-recall curves for the head-to-head comparison of CAFA2, CAFA3 and CAFA4 methods on the *limited knowledge* (LK) benchmarks

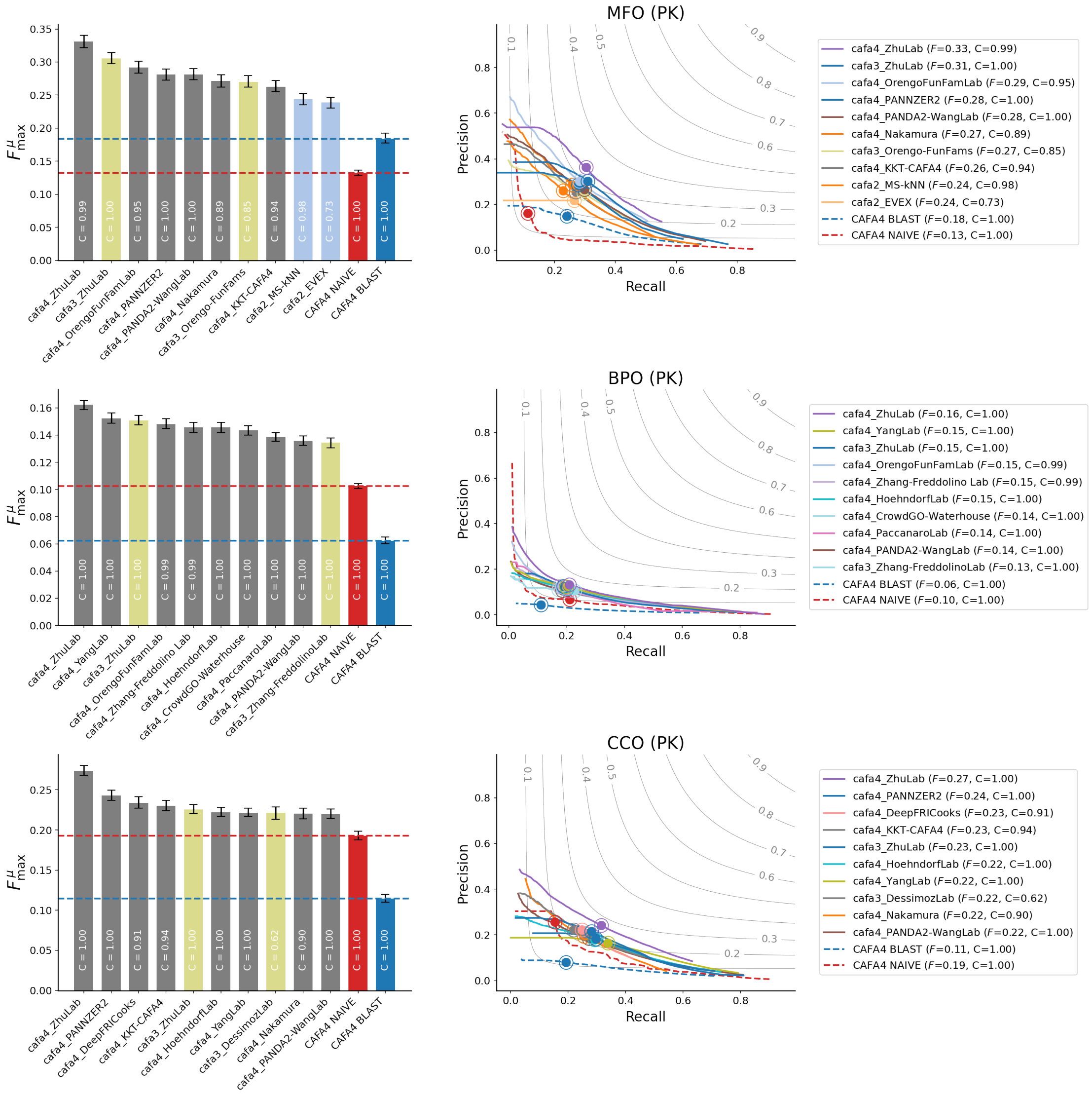


Figure S3.9: $F^{\mu}_{\max}$ scores and precision-recall curves for the head-to-head comparison of CAFA2, CAFA3 and CAFA4 methods on the *partial knowledge* (PK) benchmarks

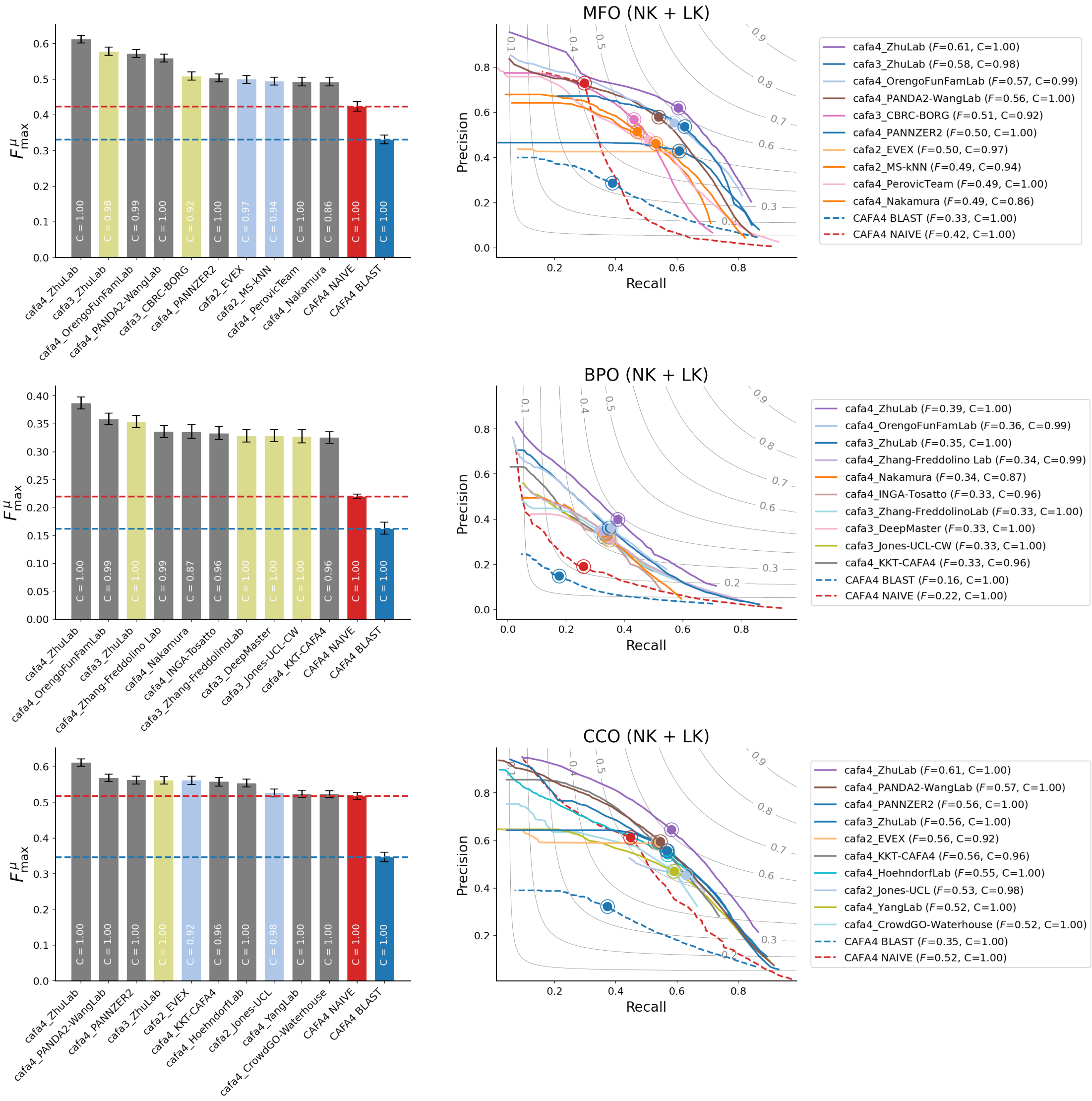


Figure S3.10: $F_{\max}^{\mu}$ scores and precision-recall curves for the head-to-head comparison of CAFA2, CAFA3 and CAFA4 methods on the *no knowledge* (NK) and *limited knowledge* (LK) benchmarks

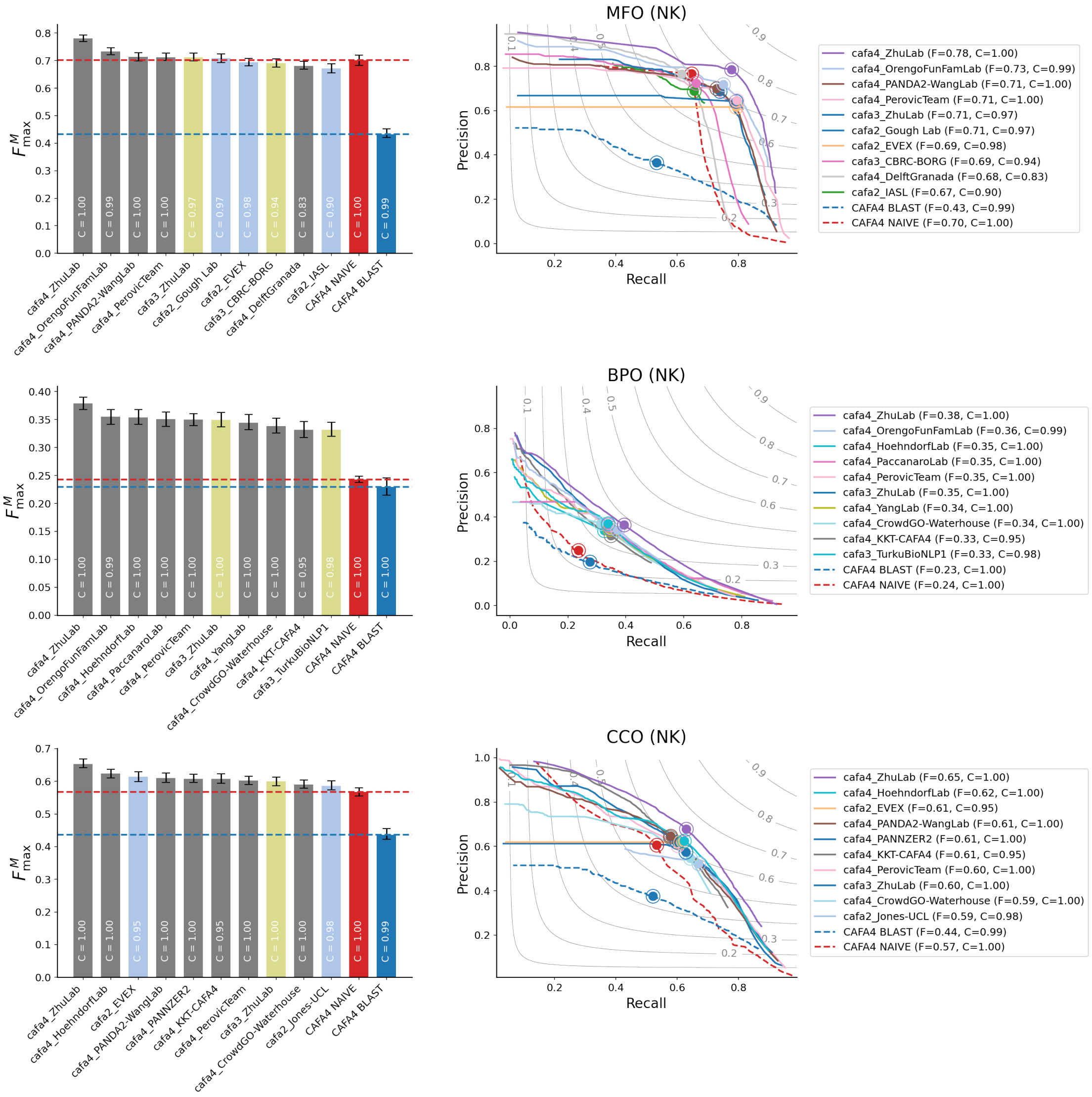

Figure S3.11: $F_{\max}^{M}$ scores and precision-recall curves for the head-to-head comparison of CAFA2, CAFA3 and CAFA4 methods on the *no knowledge* (NK) benchmarks

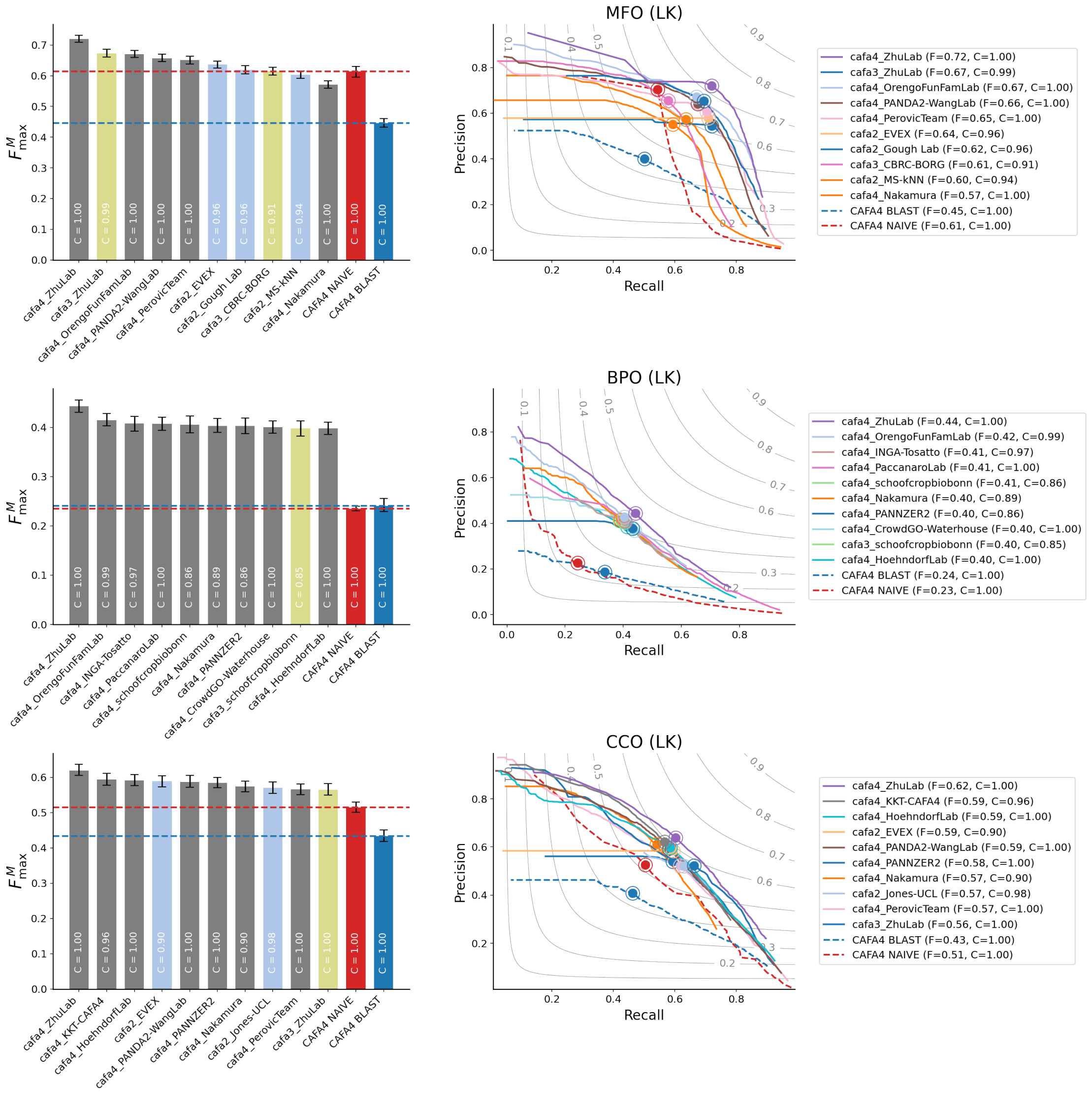


Figure S3.12: $F^M_{\text{max}}$ scores and precision-recall curves for the head-to-head comparison of CAFA2, CAFA3 and CAFA4 methods on the *limited knowledge* (LK) benchmarks

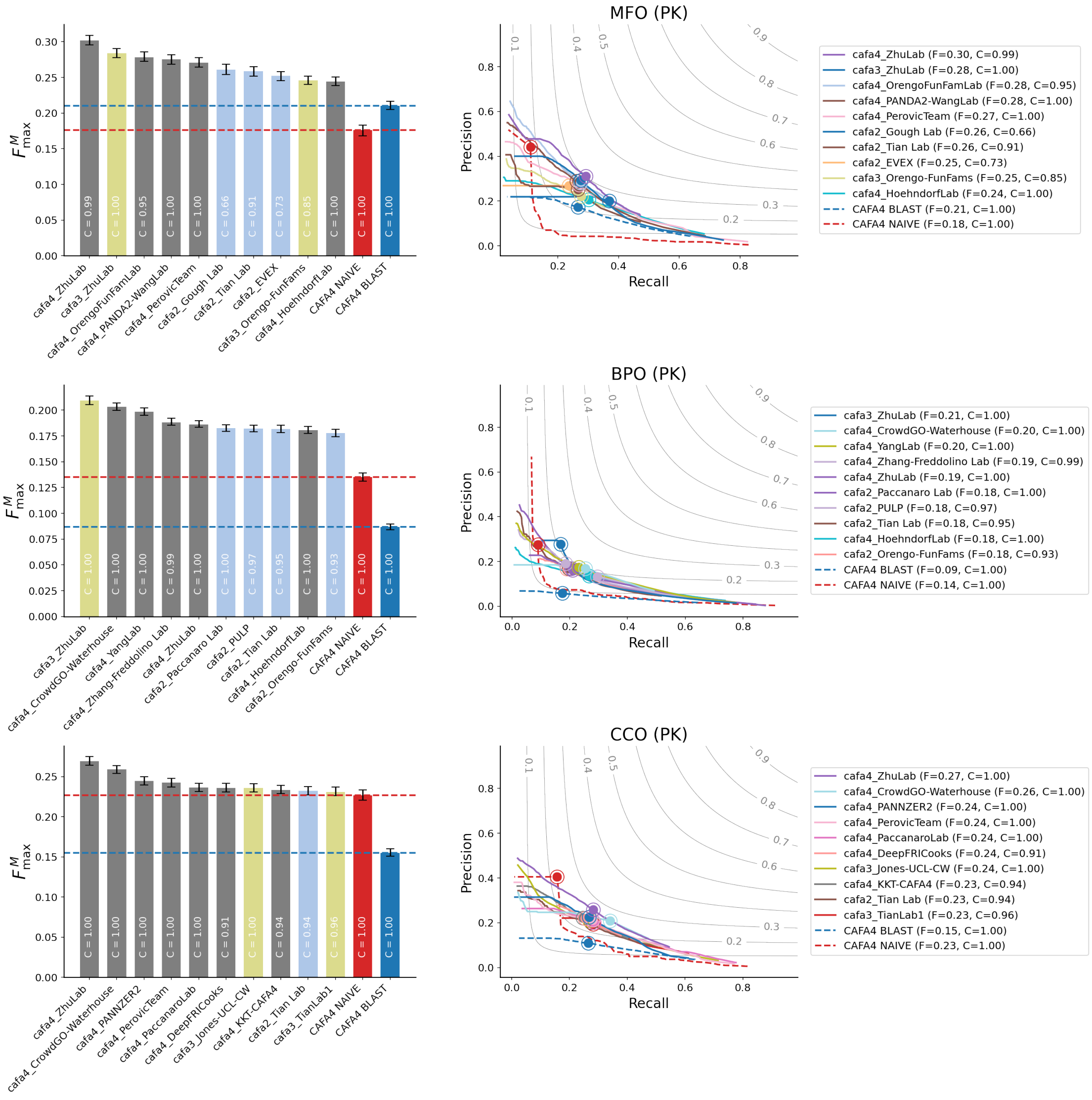


Figure S3.13: $F^{M}_{\max}$ scores and precision-recall curves for the head-to-head comparison of CAFA2, CAFA3 and CAFA4 methods on the *partial knowledge* (PK) benchmarks

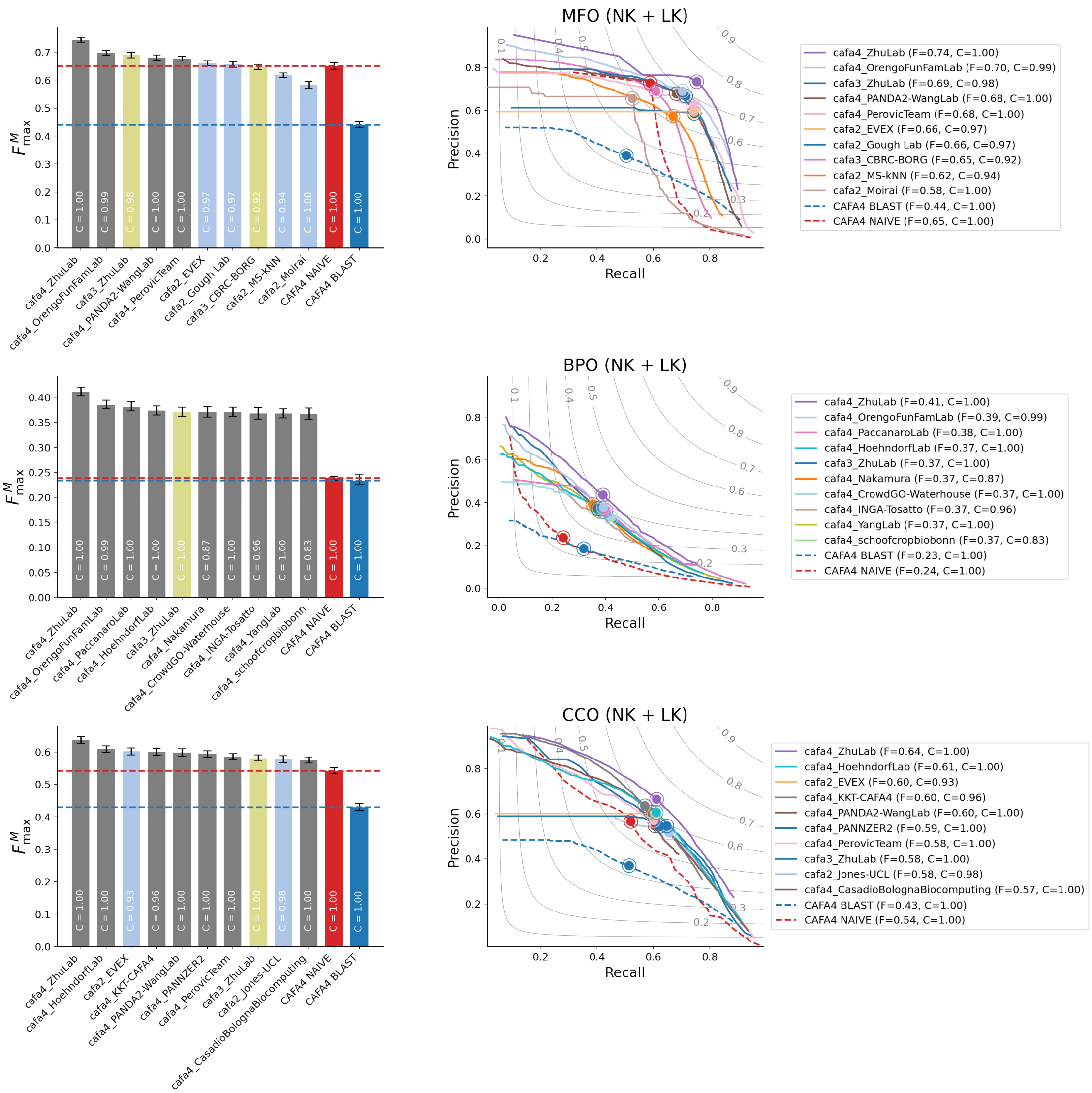


Figure S3.14: $F_{\text{max}}^{M}$ scores and precision-recall curves for the head-to-head comparison of CAFA2, CAFA3 and CAFA4 methods on the *no knowledge* (NK) and *limited knowledge* (LK) benchmarks

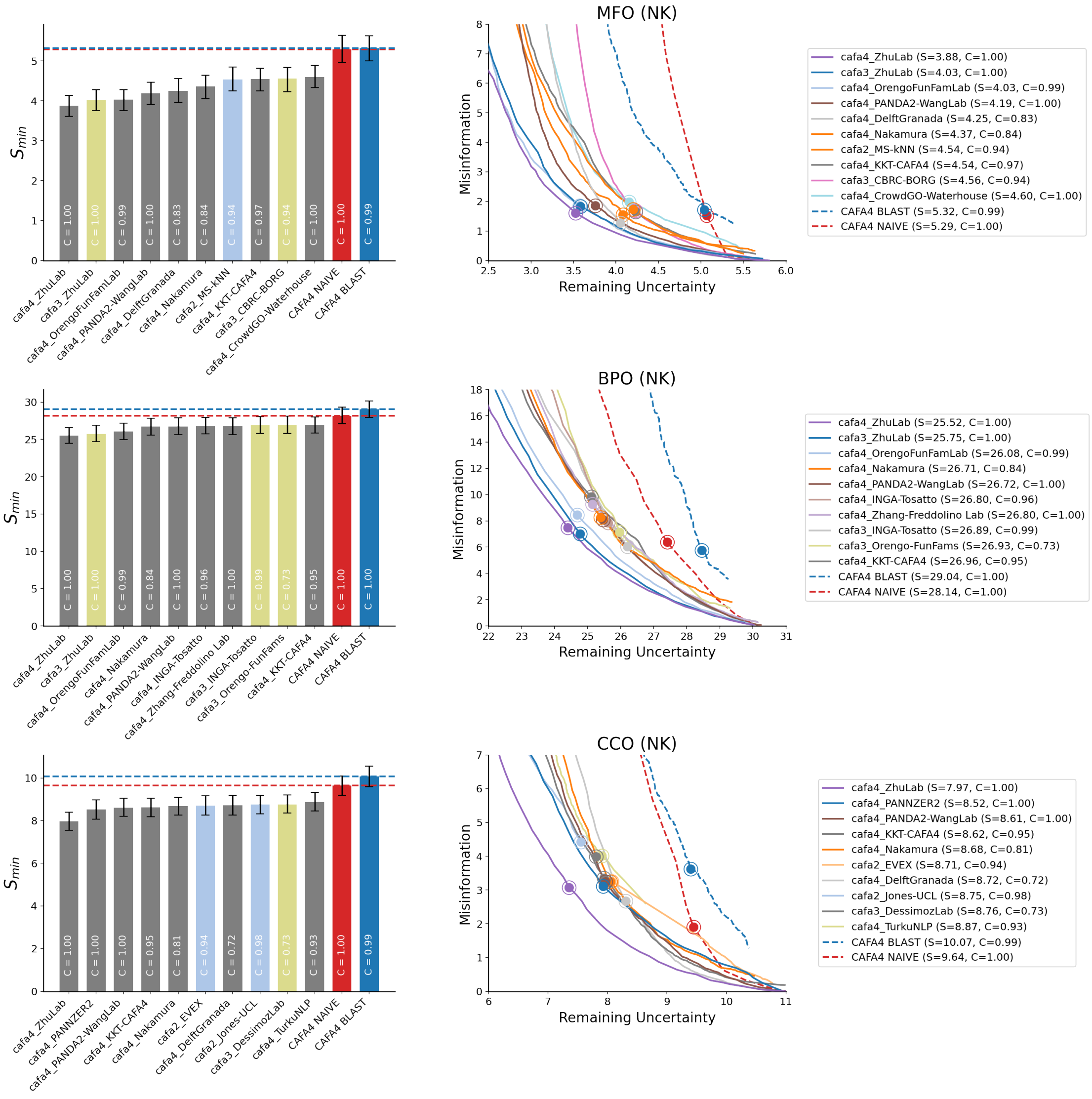


Figure S3.15: $S_{\min}$ scores and remaining uncertainty-misinformation curves for the head-to-head comparison of CAFA2, CAFA3 and CAFA4 methods on the *no knowledge* (NK) benchmarks

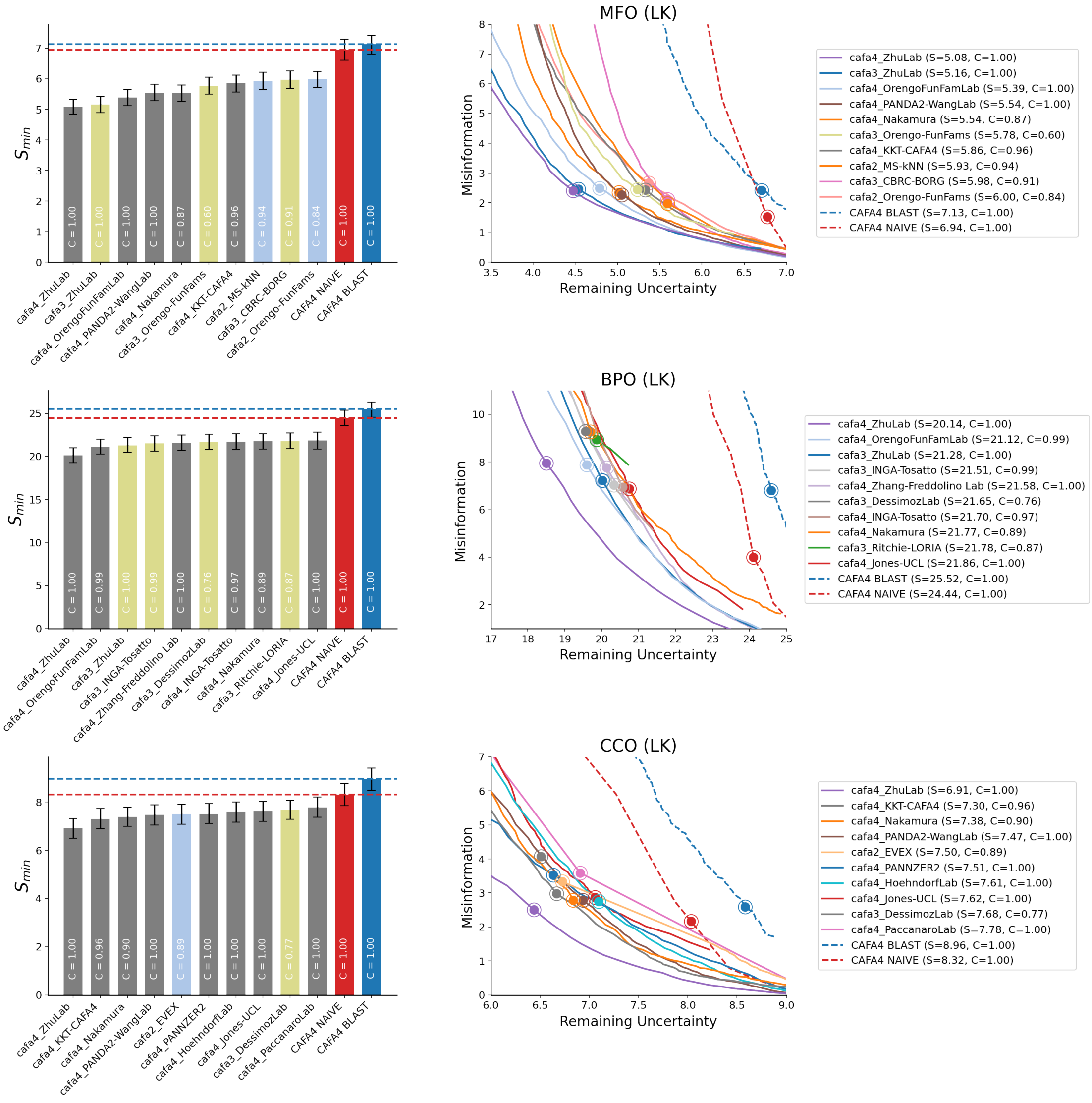


Figure S3.16: $S_{\min}$ scores and remaining uncertainty-misinformation curves for the head-to-head comparison of CAFA2, CAFA3 and CAFA4 methods on the *limited knowledge* (LK) benchmarks

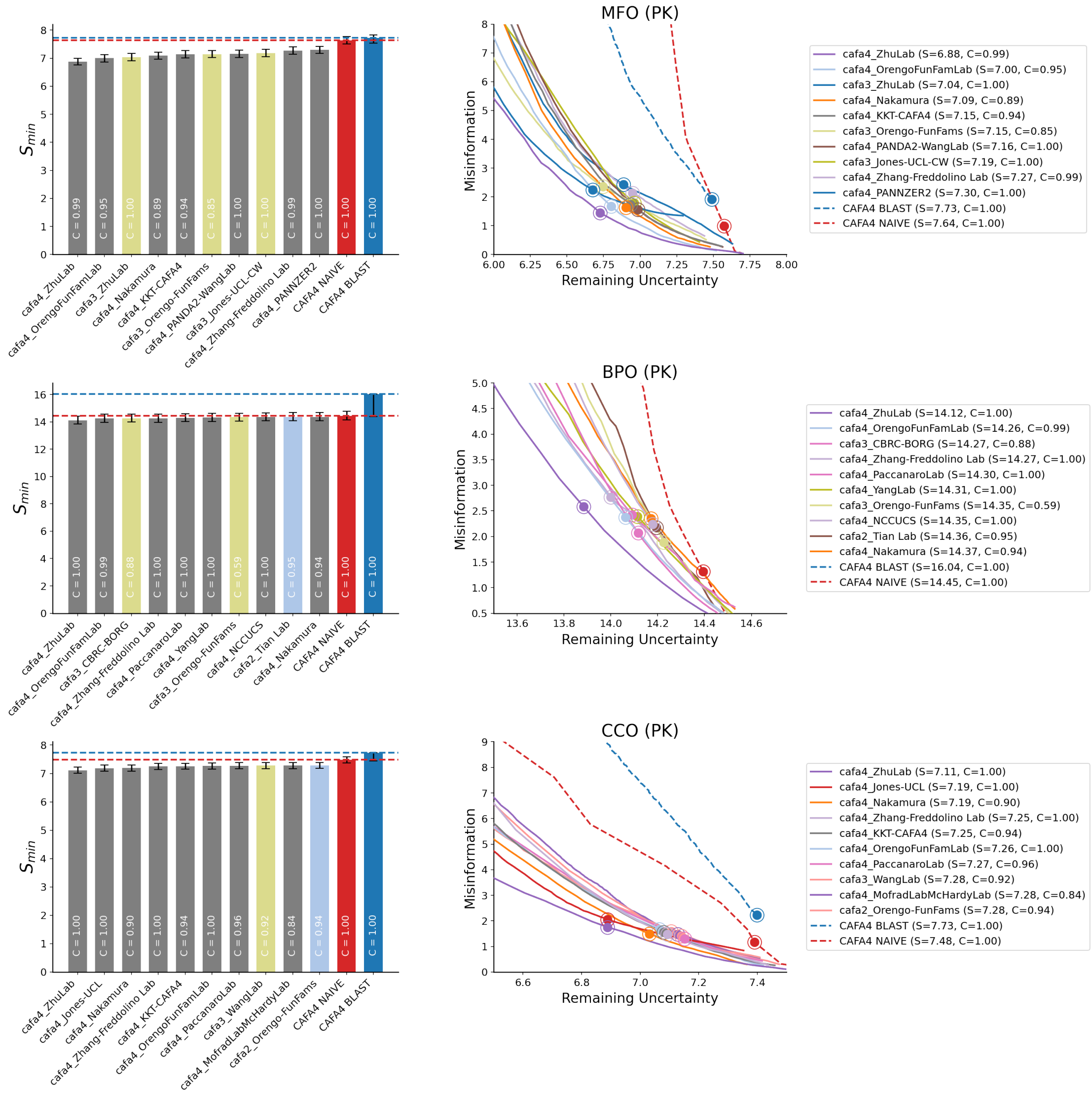


Figure S3.17: $S_{\min}$ scores and remaining uncertainty-misinformation curves for the head-to-head comparison of CAFA2, CAFA3 and CAFA4 methods on the *partial knowledge* (PK) benchmarks

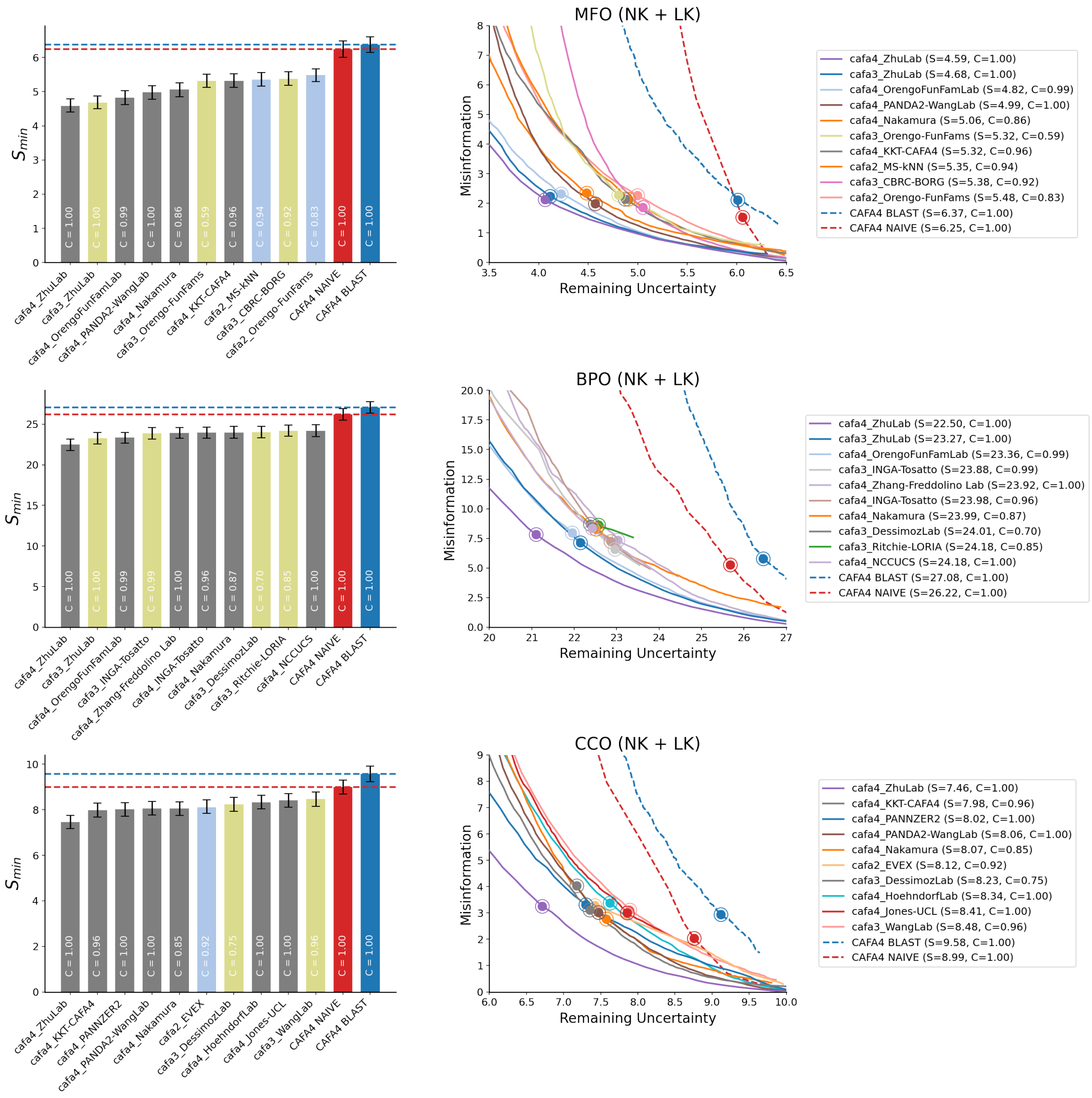


Figure S3.18: $S_{\text{min}}$ scores and remaining uncertainty-misinformation curves for the head-to-head comparison of CAFA2, CAFA3 and CAFA4 methods on combined *no knowledge* (NK) and *limited knowledge* (LK) benchmarks

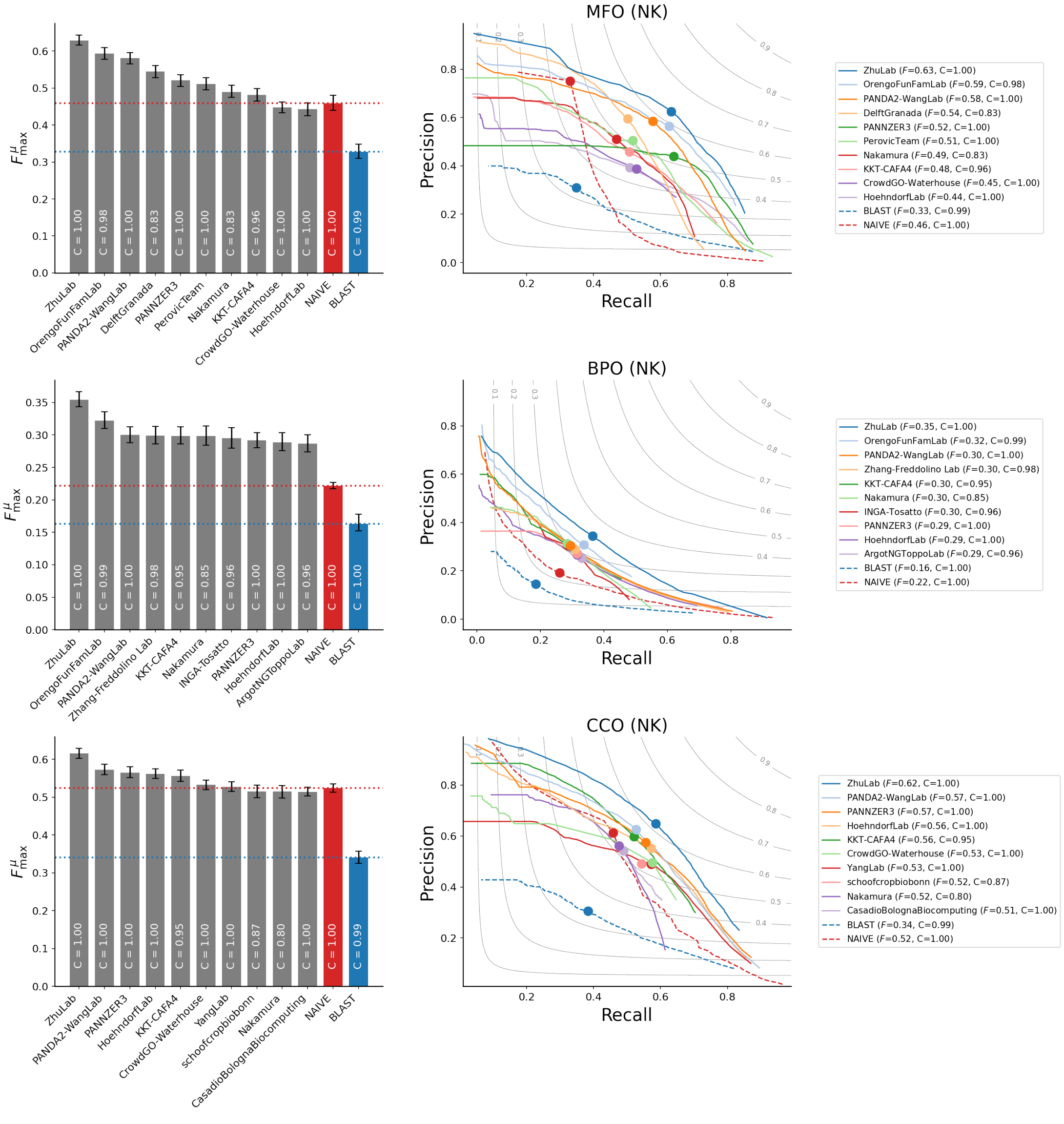


Figure S3.19: $F^{\mu}_{\max}$ scores and precision-recall curves for CAFA4 methods, NAIVE and BLAST baselines on the *no knowledge* (NK) benchmarks

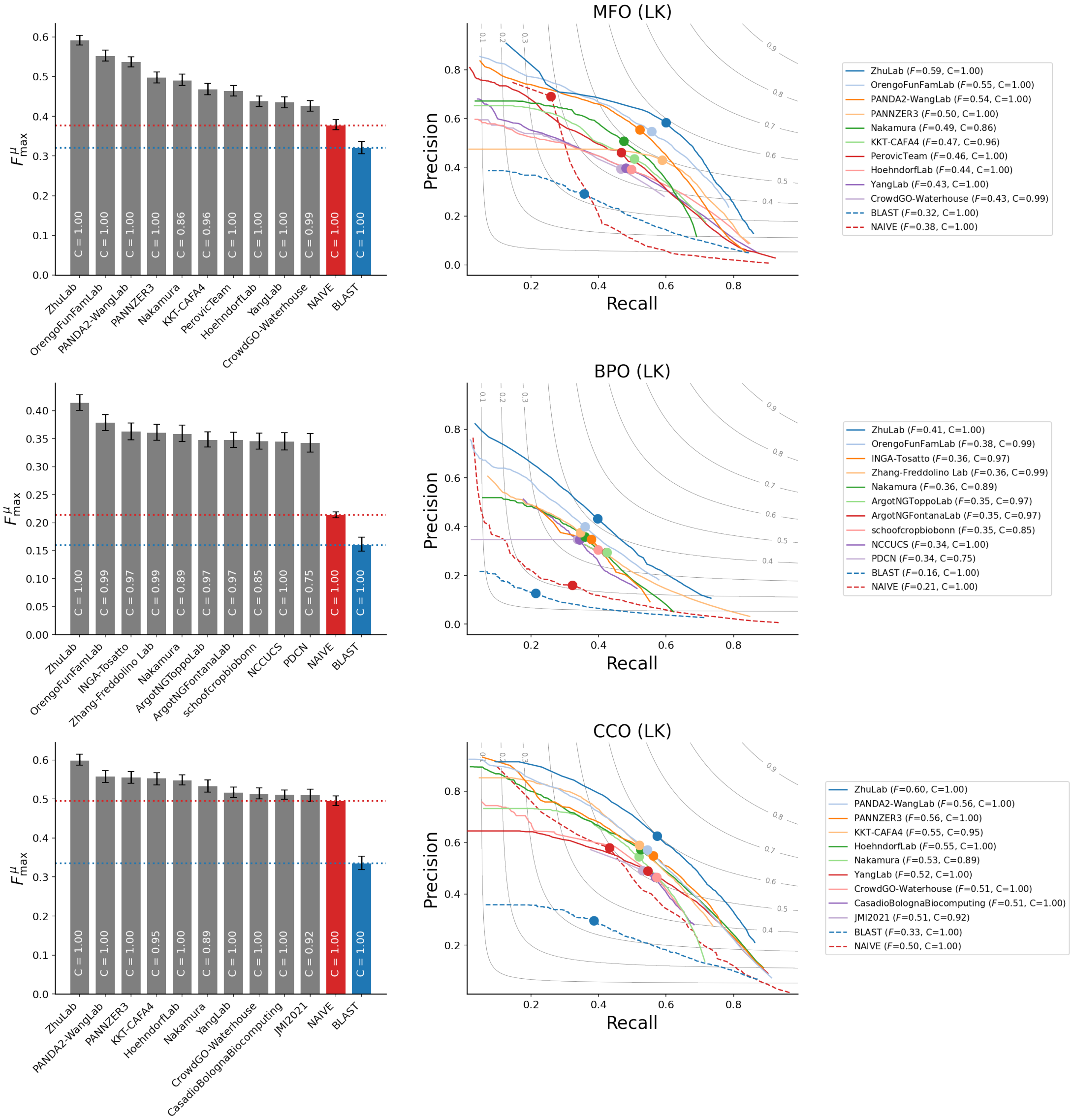


Figure S3.20: $F_{max}^{\mu}$ scores and precision-recall curves for CAFA4 methods, NAIVE and BLAST baselines on the *limited knowledge* (LK) benchmarks

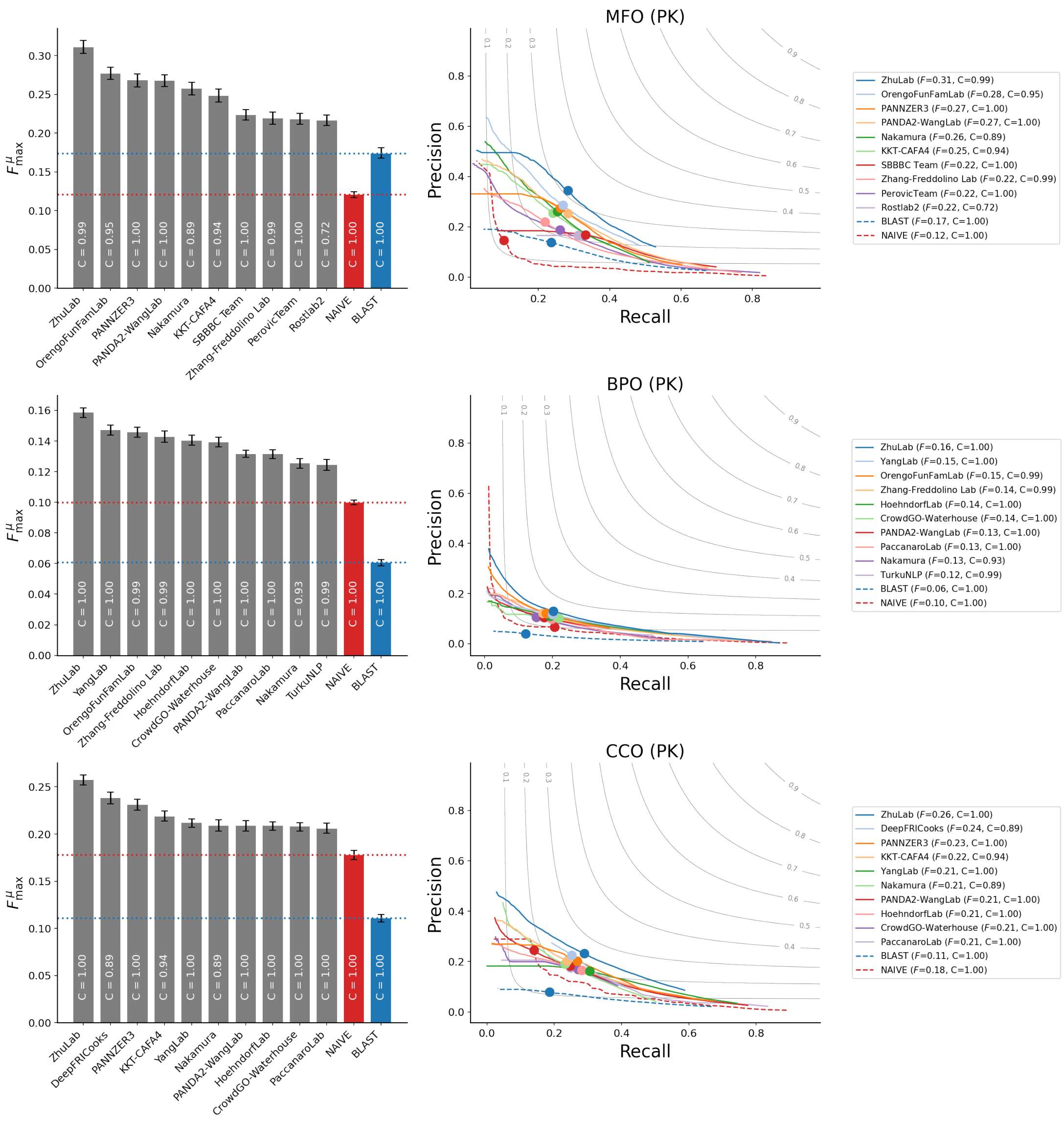


Figure S3.21: $F_{\max}^{\mu}$ scores and precision-recall curves for CAFA4 methods, NAIVE and BLAST baselines on the *partial knowledge* (PK) benchmarks

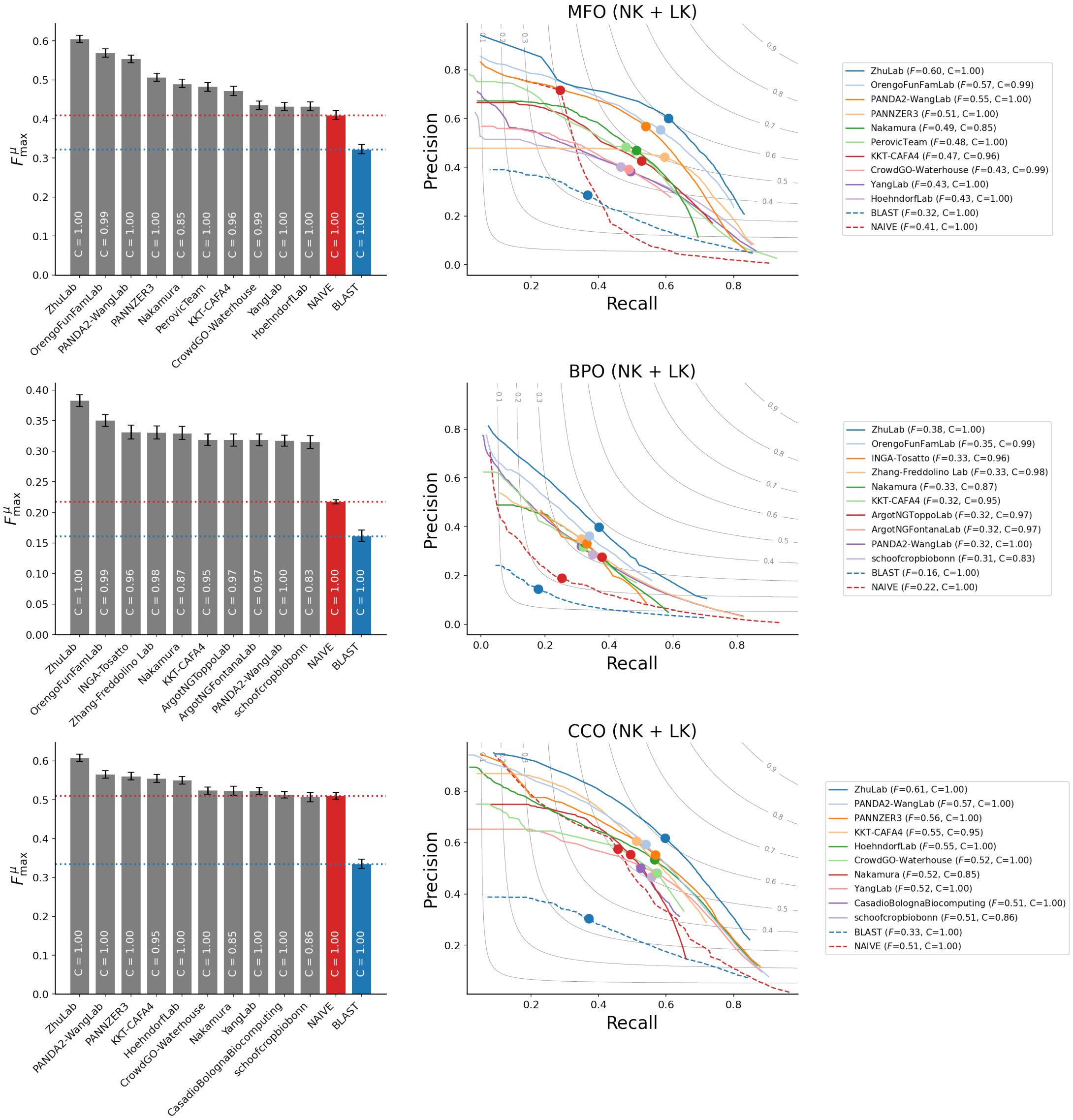


Figure S3.22: $F^{\mu}_{\max}$ scores and precision-recall curves for CAFA4 methods, NAIVE and BLAST baselines on combined *no knowledge* (NK) and *limited knowledge* (LK) benchmarks

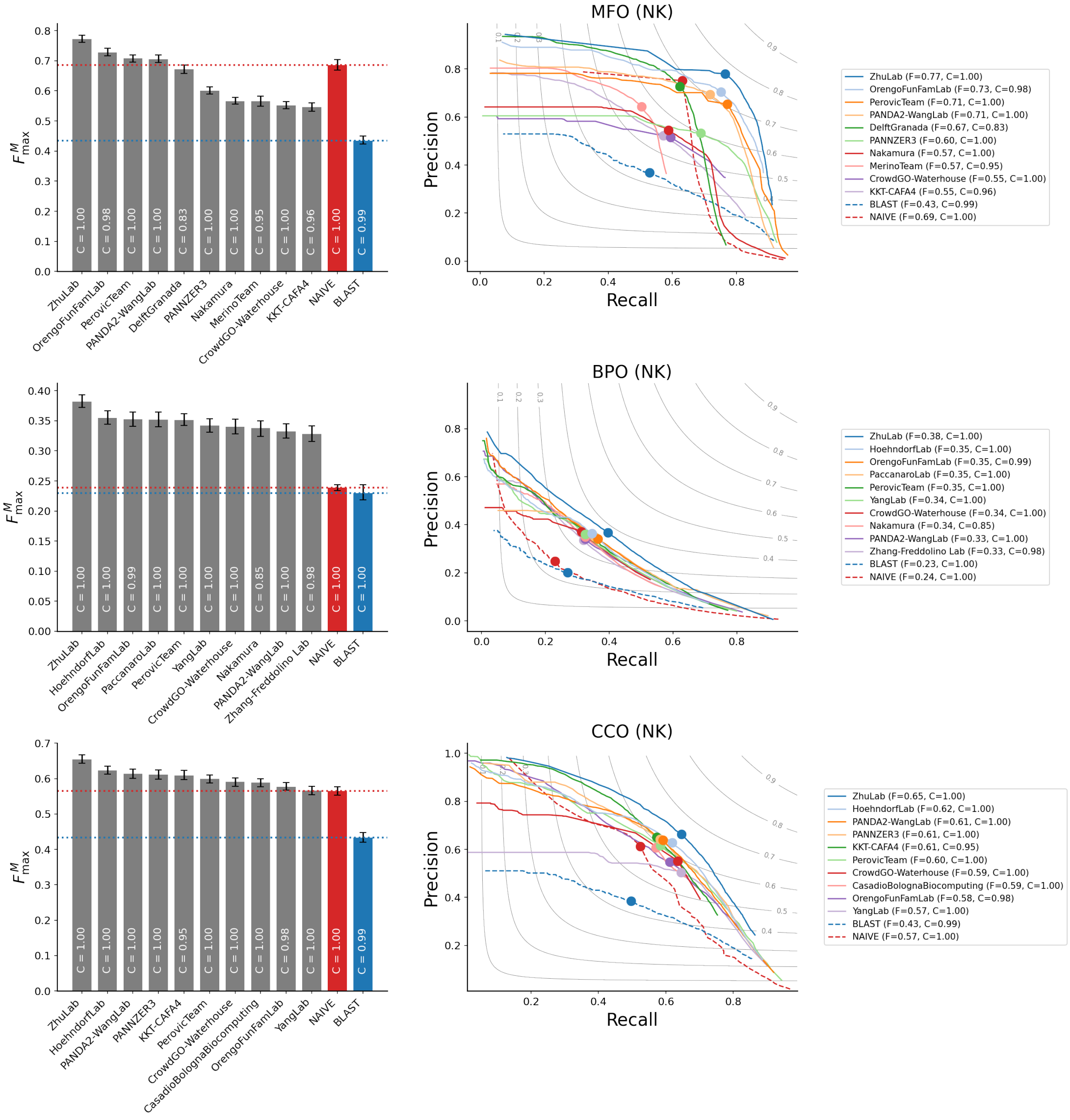


Figure S3.23: $F_{\text{max}}^{M}$ scores and precision-recall curves for CAFA4 methods, NAIVE and BLAST baselines on the *no knowledge* (NK) benchmarks

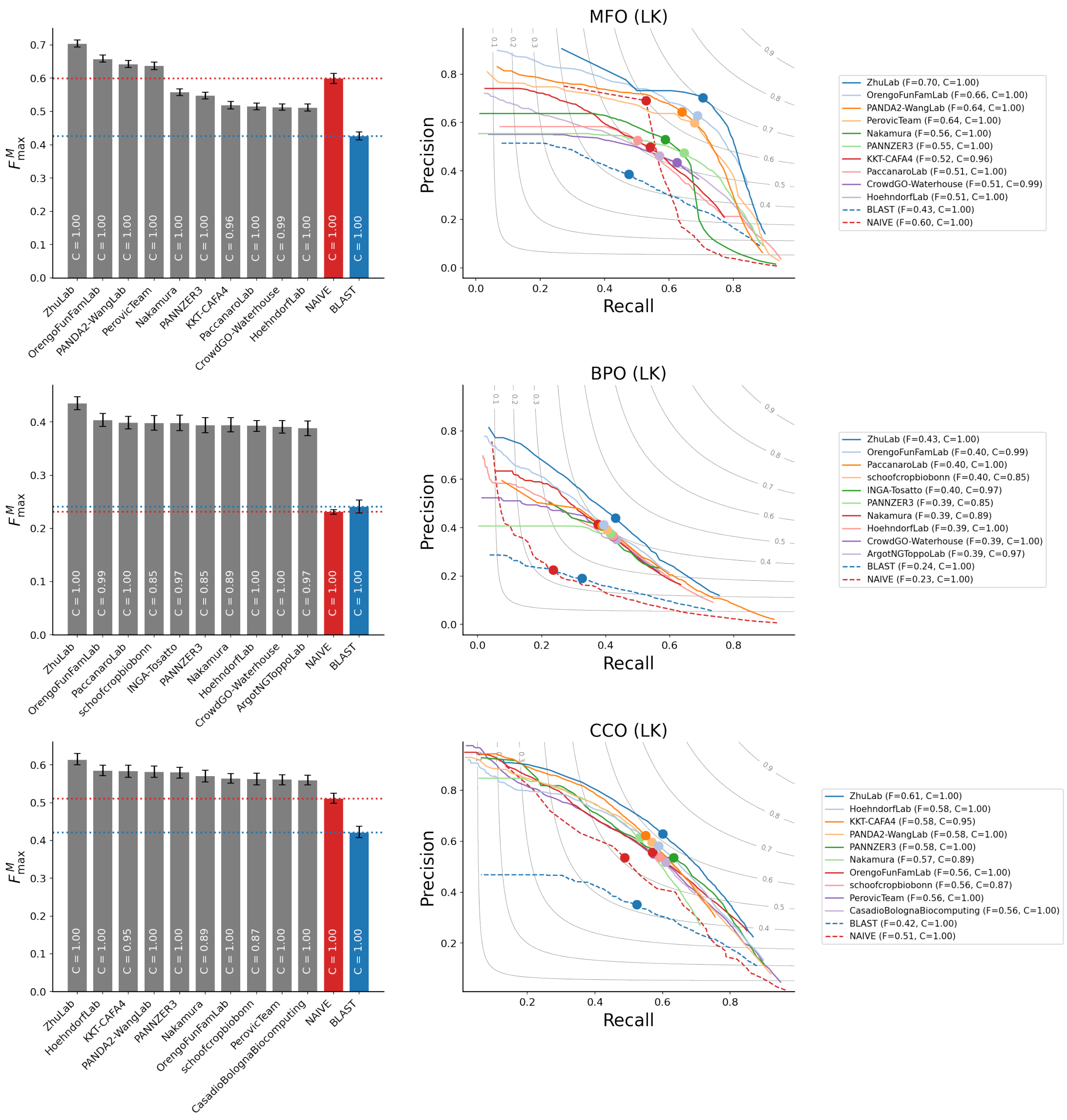


Figure S3.24: $F^M_{\max}$ scores and precision-recall curves for CAFA4 methods, NAIVE and BLAST baselines on the *limited knowledge* (LK) benchmarks

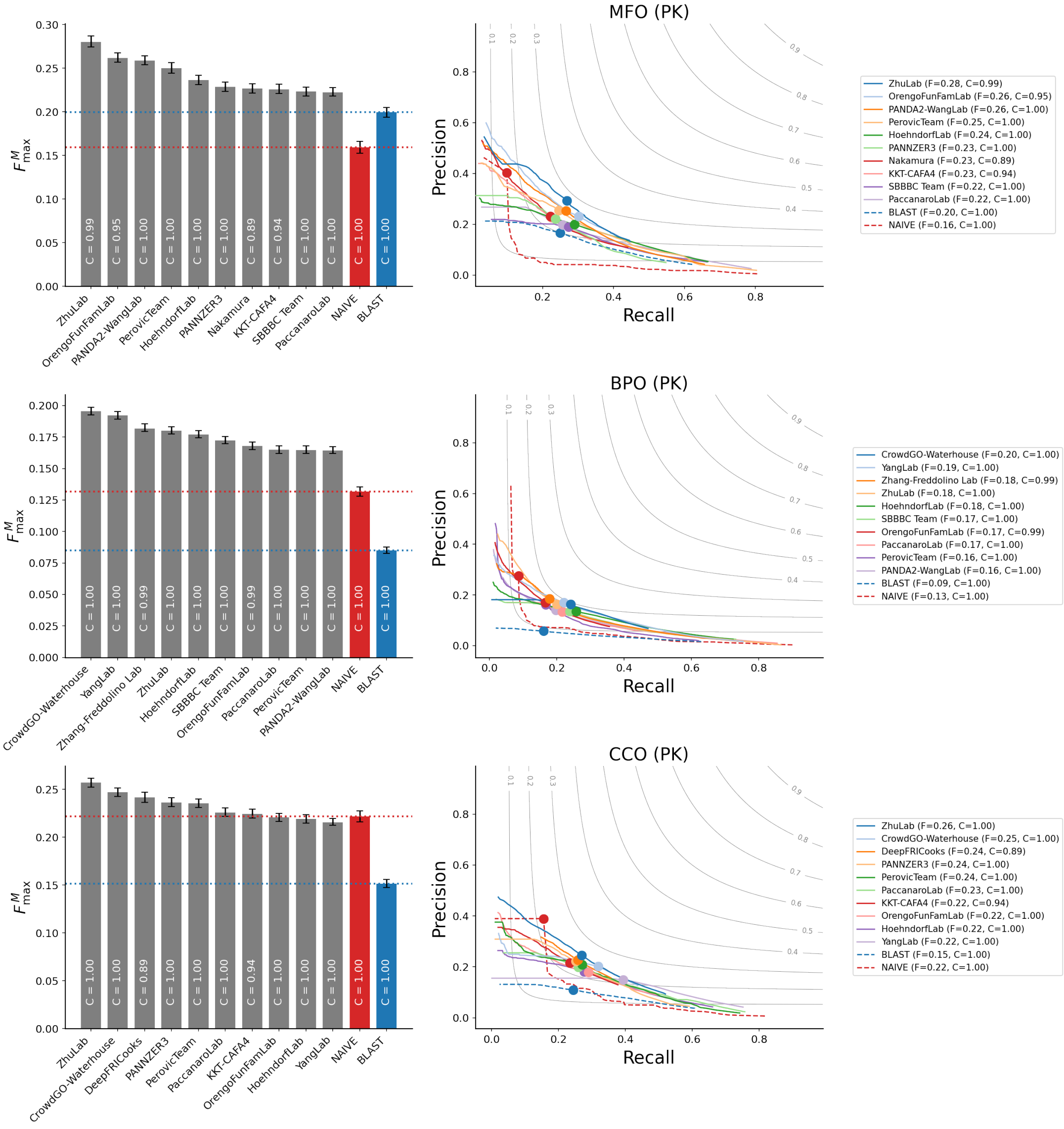


Figure S3.25: $F_{\max}^{M}$ scores and precision-recall curves for CAFA4 methods, NAIVE and BLAST baselines on the *partial knowledge* (PK) benchmarks

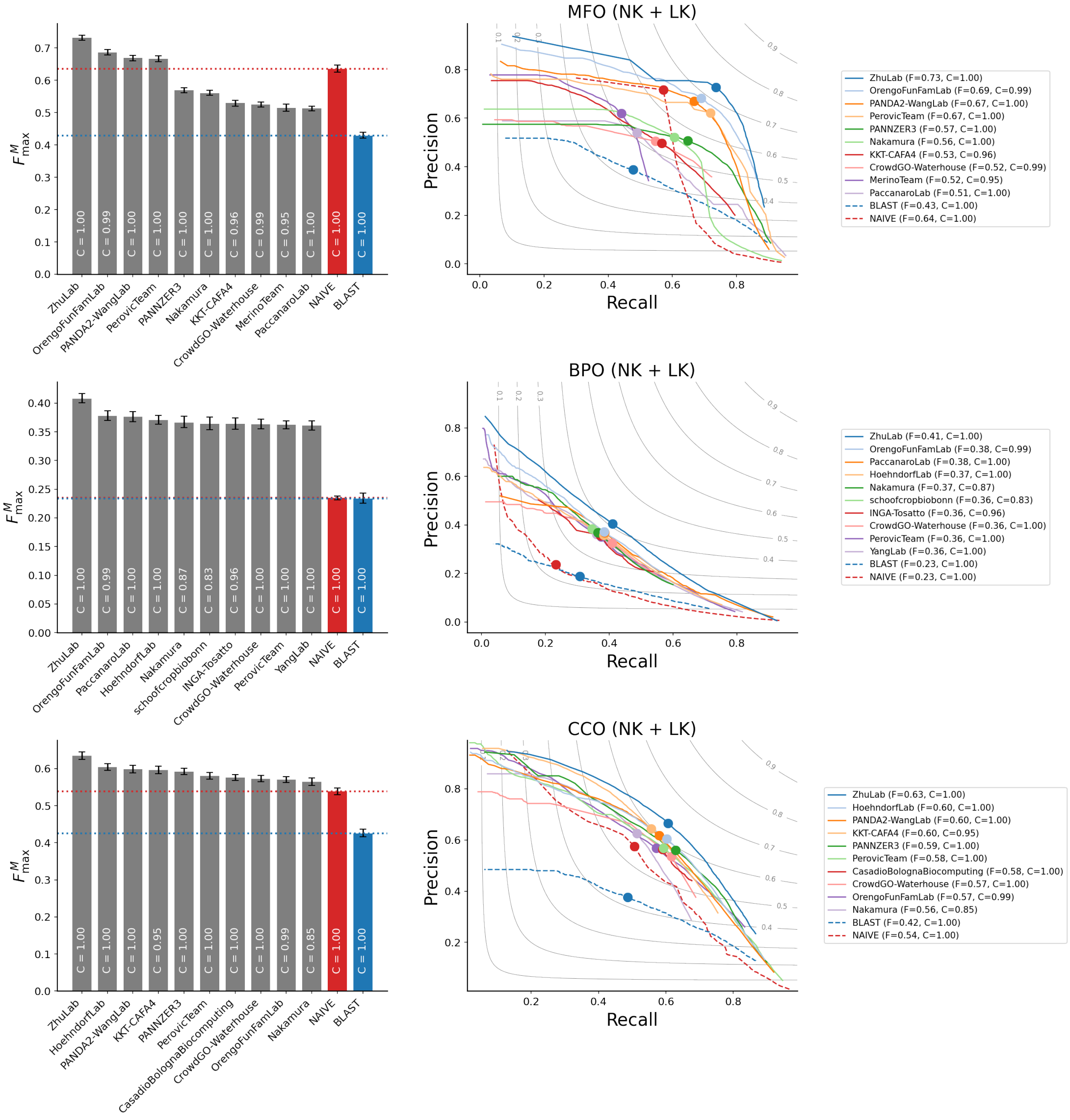


Figure S3.26: $F_{\max}^{M}$ scores and precision-recall curves for CAFA4 methods, NAIVE and BLAST baselines on combined *no knowledge* (NK) and *limited knowledge* (LK) benchmarks

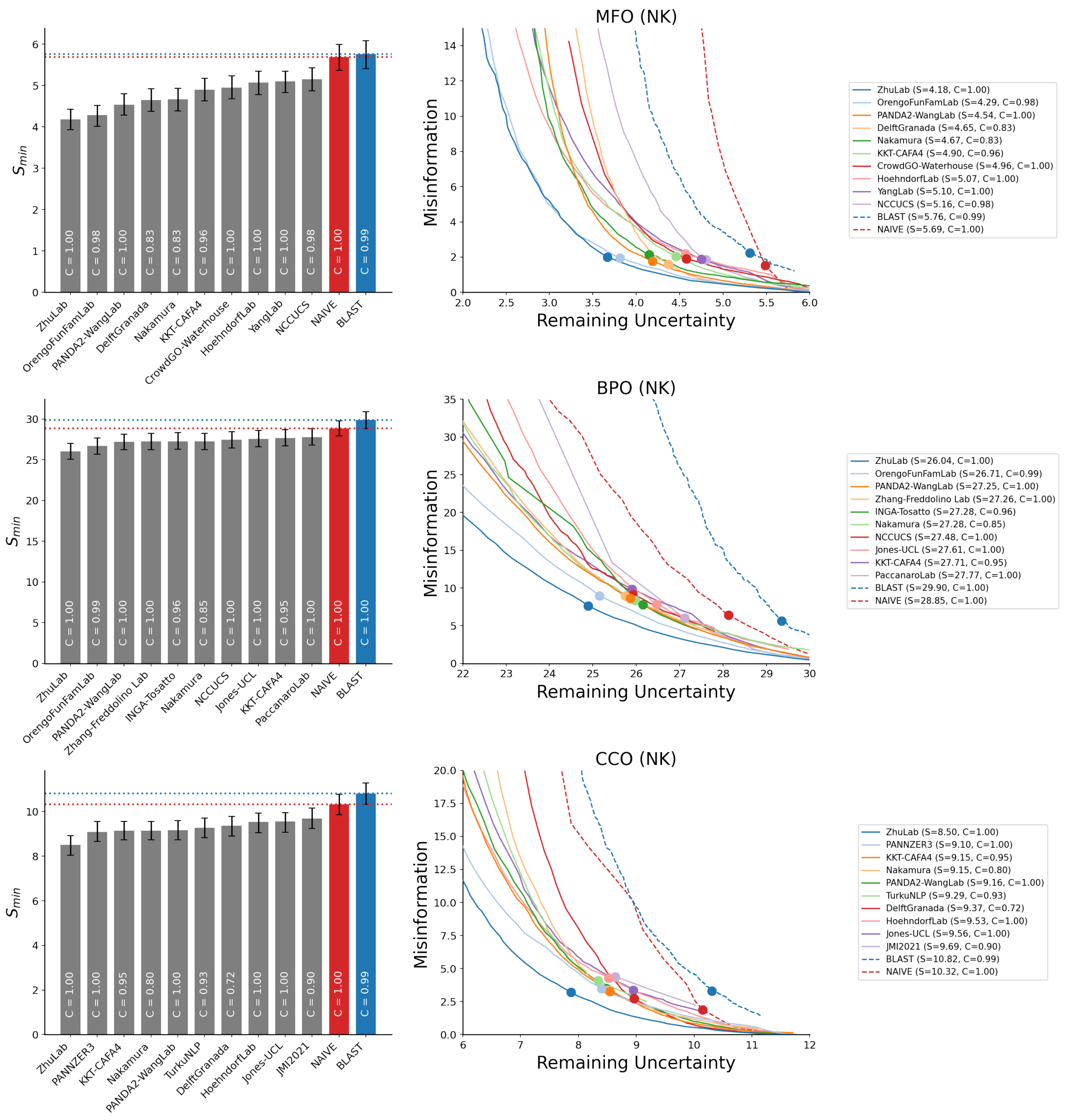


Figure S3.27: $S_{\min}$ scores and remaining uncertainty-misinformation curves for CAFA4 methods, NAIVE and BLAST baselines on the *no knowledge* (NK) benchmarks

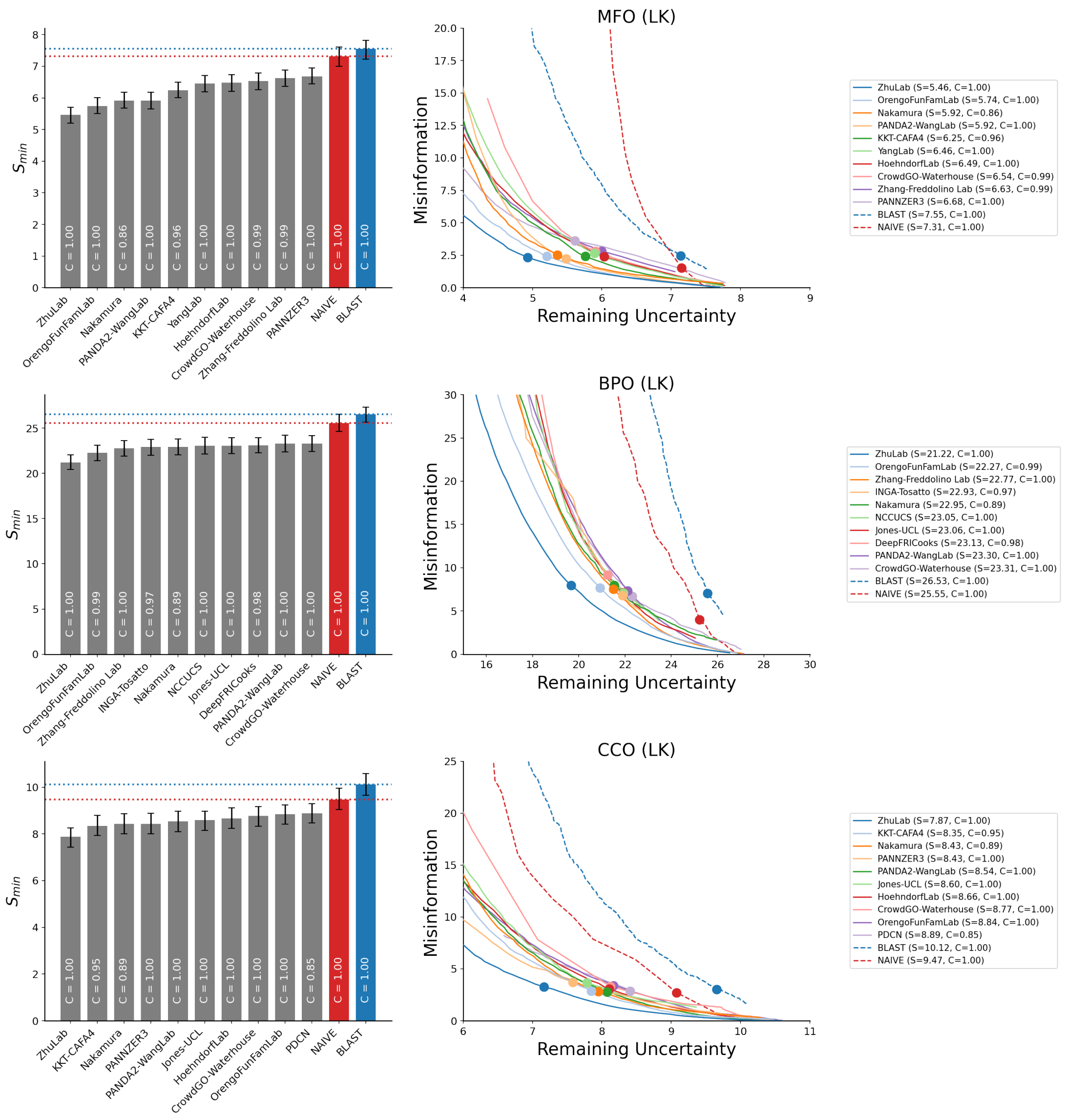


Figure S3.28: $S_{\text{min}}$ scores and remaining uncertainty-misinformation curves for CAFA4 methods, NAIVE and BLAST baselines on the *limited knowledge* (LK) benchmarks

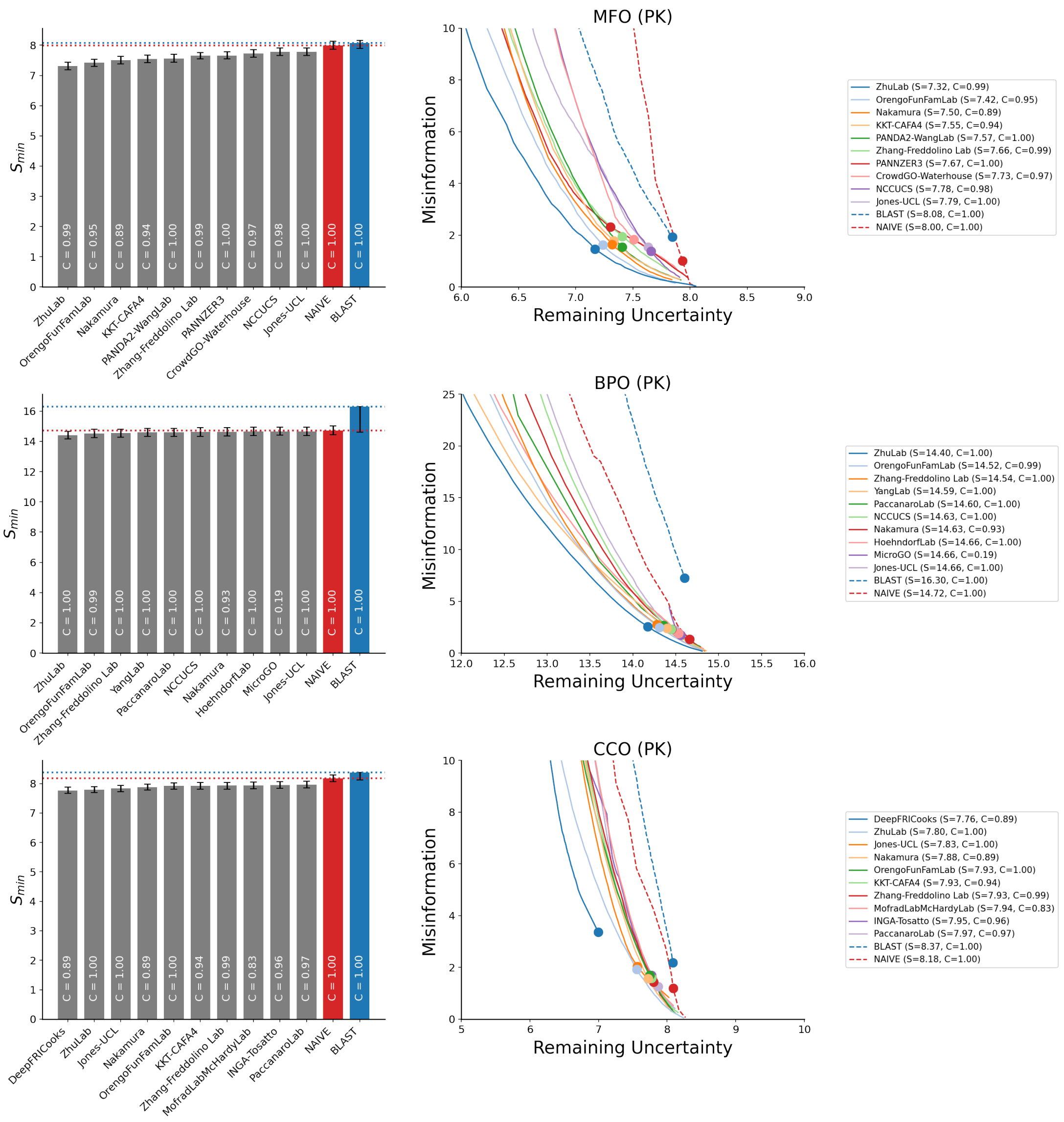


Figure S3.29: $S_{\min}$ scores and remaining uncertainty-misinformation curves for CAFA4 methods, NAIVE and BLAST baselines on the *partial knowledge* (PK) benchmarks

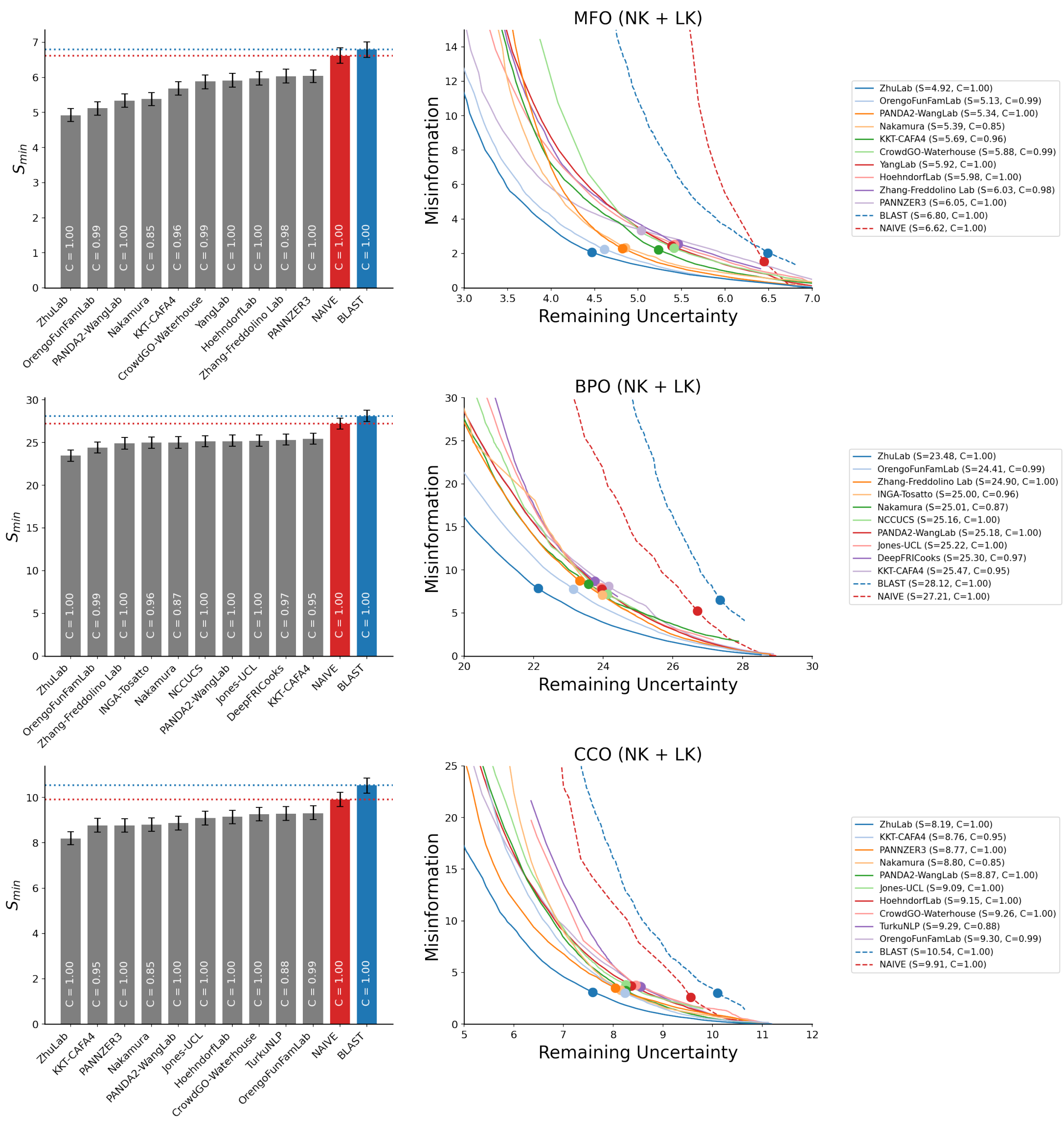


Figure S3.30: $S_{\min}$ scores and remaining uncertainty-misinformation curves for CAFA4 methods, NAIVE and BLAST baselines on combined *no knowledge* (NK) and *limited knowledge* (LK) benchmarks

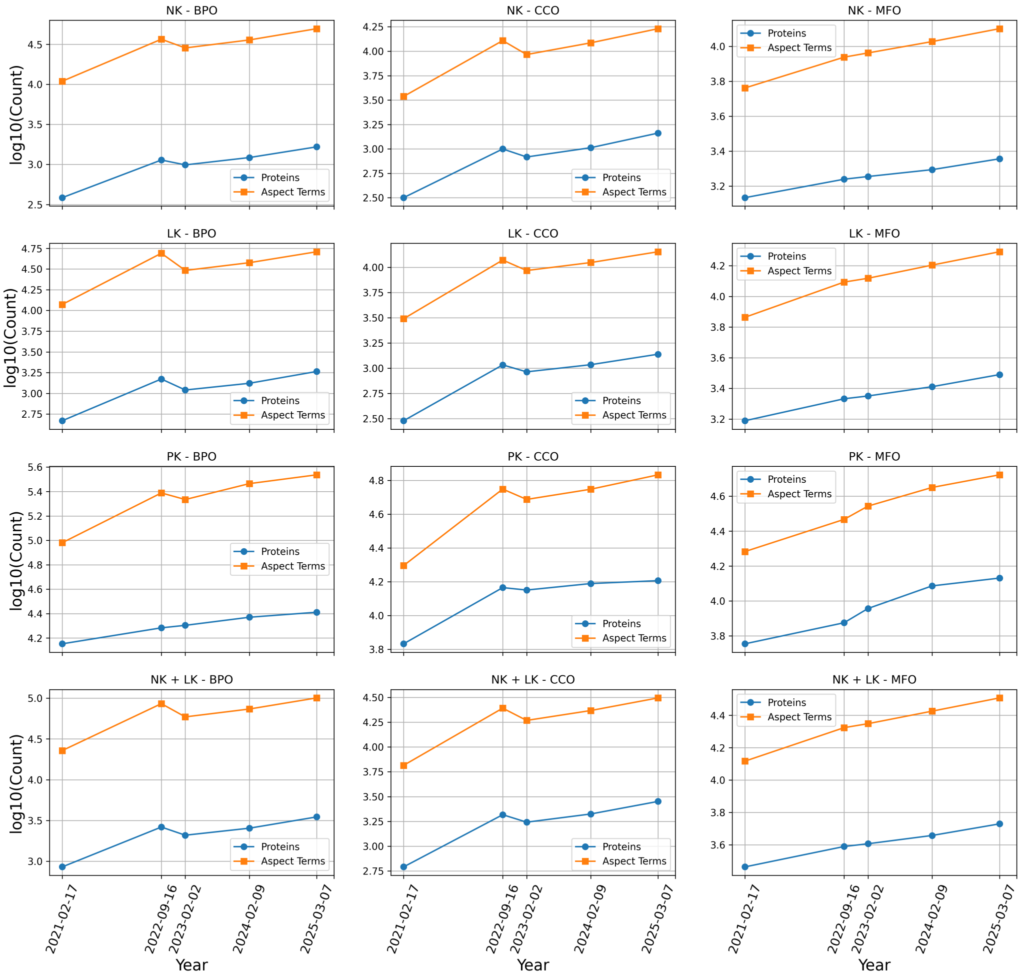


Figure S3.31: Growth in numbers of proteins and gene ontology (GO) annotations in the benchmark datasets. Number of proteins and gene ontology (GO) annotations at $\mathcal{B}_{21}$, $\mathcal{B}_{22}$, $\mathcal{B}_{23}$, $\mathcal{B}_{24}$ and $\mathcal{B}_{25}$ for *no knowledge* (NK), *limited knowledge* (LK), *partial knowledge* (PK) and combined *no knowledge* (NK) and *limited knowledge* (LK) benchmarks in three GO aspects - Molecular Function ontology (MFO), Biological Process ontology (BPO), Cellular Component Ontology (CCO) on log scale

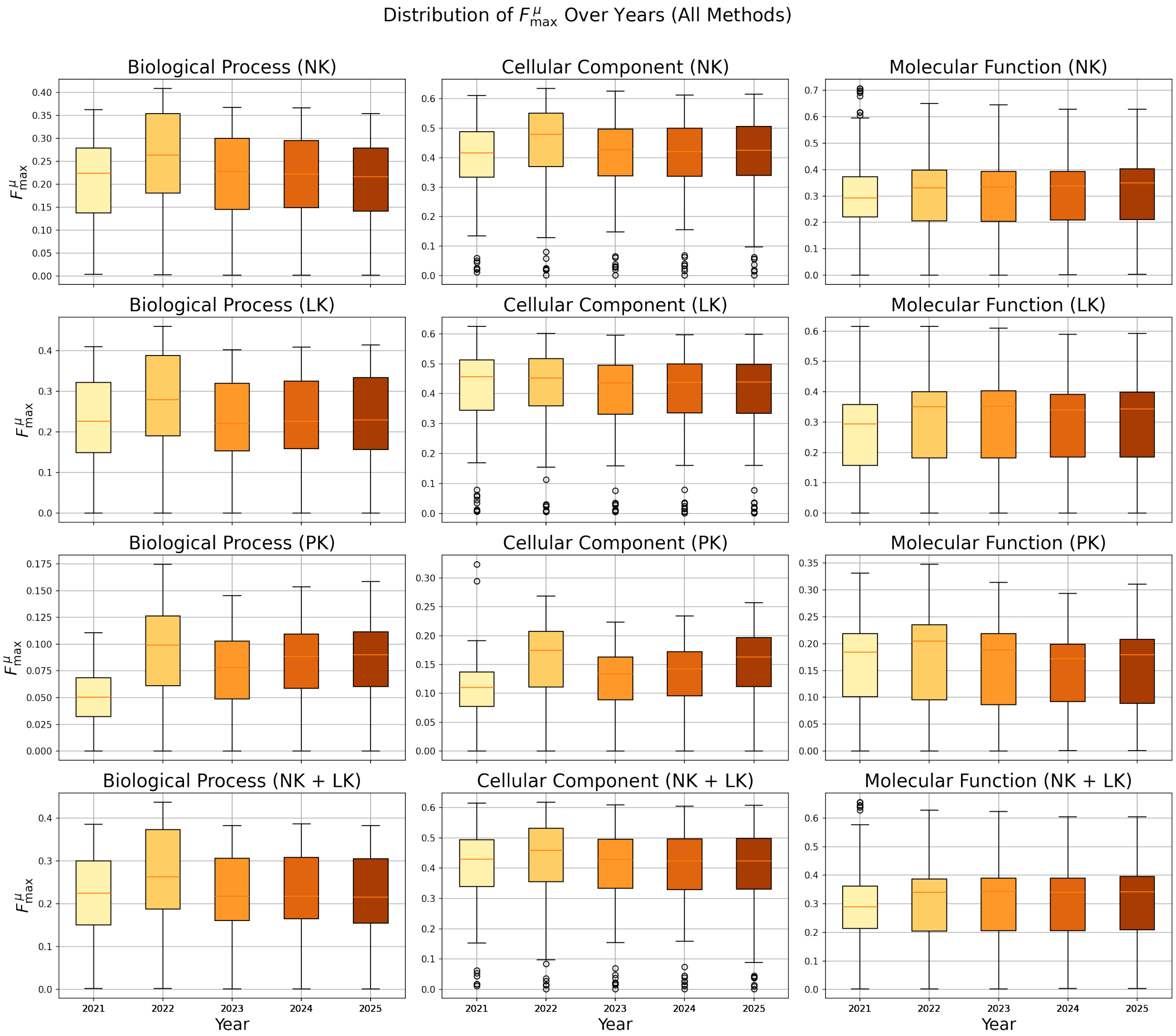


Figure S3.32: The distributions of the $F^{\mu}_{\max}$ of all CAFA4 methods calculated on the benchmarks at $\mathcal{B}_{21}$, $\mathcal{B}_{22}$, $\mathcal{B}_{23}$, $\mathcal{B}_{24}$ and $\mathcal{B}_{25}$ for *no knowledge* (NK), *limited knowledge* (LK), *partial knowledge* (PK) and combined *no knowledge* (NK) and *limited knowledge* (LK) in three GO aspects - Molecular Function ontology (MFO), Biological Process ontology (BPO), Cellular Component Ontology (CCO)

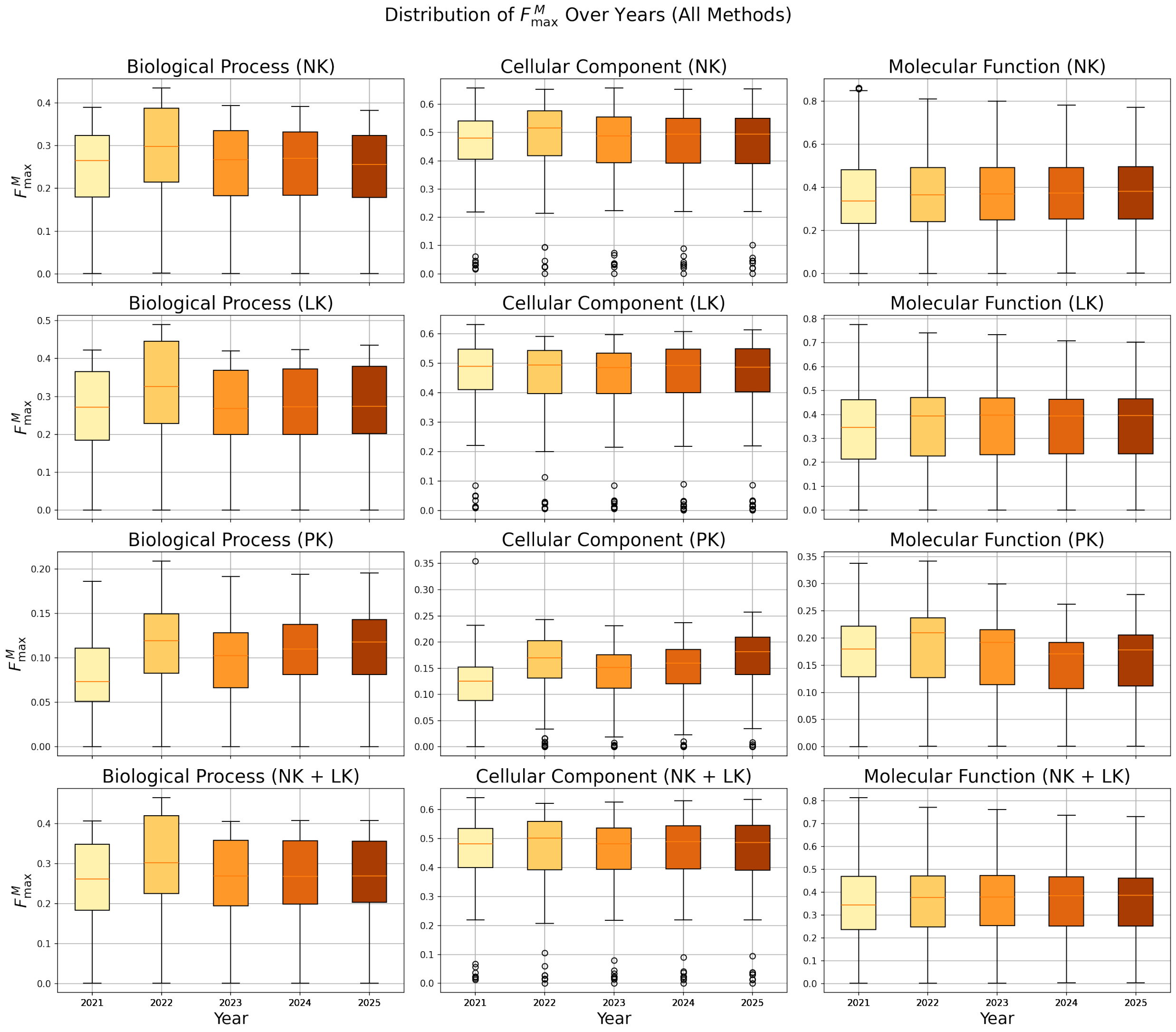


Figure S3.33: The distributions of the $F_{\max}^M$ of all CAFA4 methods calculated on the benchmarks at $\mathcal{B}_{21}$, $\mathcal{B}_{22}$, $\mathcal{B}_{23}$, $\mathcal{B}_{24}$ and $\mathcal{B}_{25}$ for *no knowledge* (NK), *limited knowledge* (LK), *partial knowledge* (PK) and combined *no knowledge* (NK) and *limited knowledge* (LK) in three GO aspects - Molecular Function ontology (MFO), Biological Process ontology (BPO), Cellular Component Ontology (CCO)

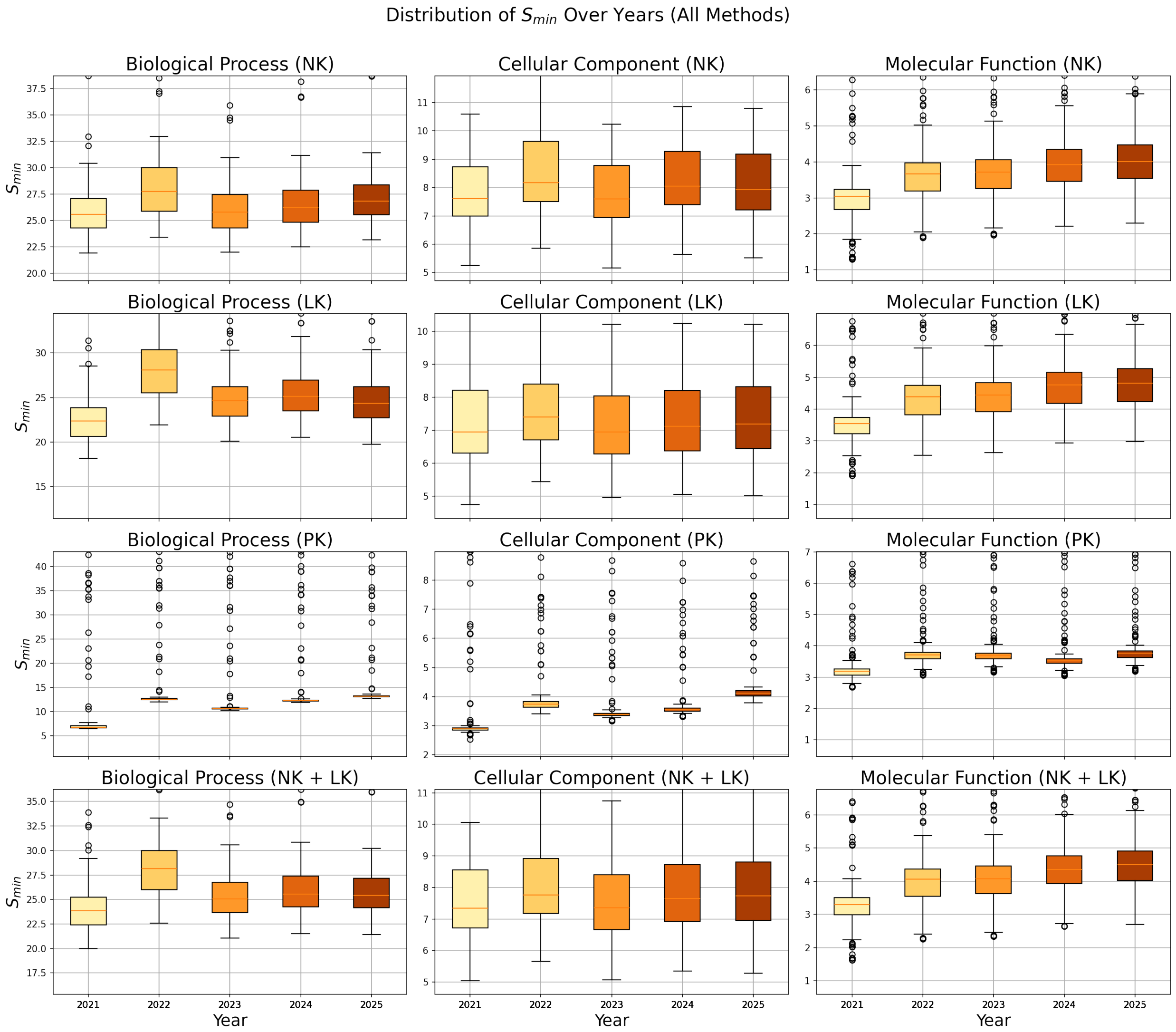


Figure S3.34: The distributions of the $S_{\text{min}}$ of all CAFA4 methods calculated on the benchmarks at $\mathcal{B}_{21}$, $\mathcal{B}_{22}$, $\mathcal{B}_{23}$, $\mathcal{B}_{24}$ and $\mathcal{B}_{25}$ for *no knowledge* (NK), *limited knowledge* (LK), *partial knowledge* (PK) and combined *no knowledge* (NK) and *limited knowledge* (LK) in three GO aspects - Molecular Function ontology (MFO), Biological Process ontology (BPO), Cellular Component Ontology (CCO)

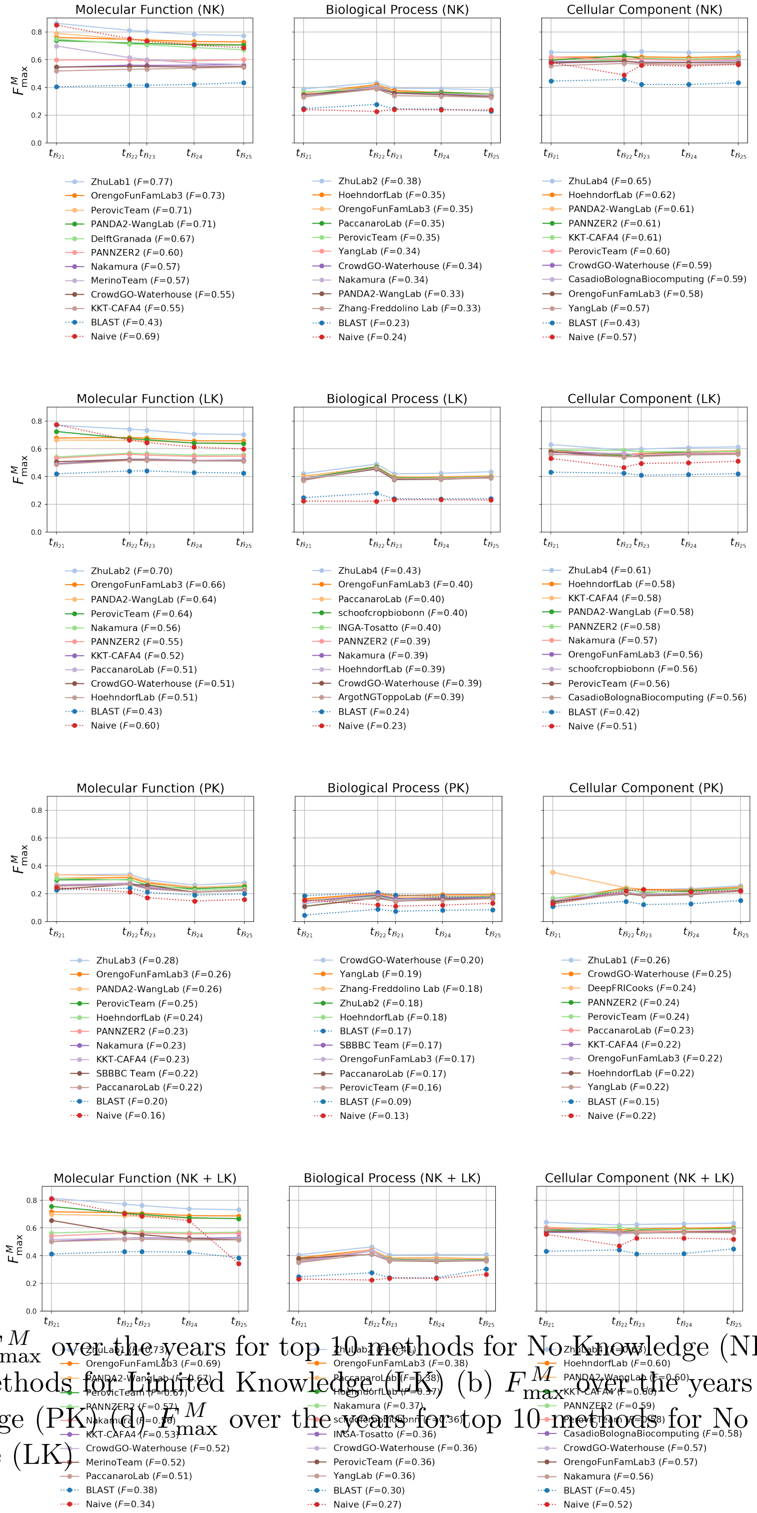


Figure S3.35: (a) $F^M_{\max}$ over the years for top 10 methods for No Knowledge (NK) (b) $F^M_{\max}$over the years for top 10 methods for Limited Knowledge (LK) (b) $F^M_{\max}$ over the years for top 10 methods for partial knowledge (PK) (d) $F^M_{\max}$ over the years for top 10 methods for No Knowledge (NK) + Limited Knowledge (LK)

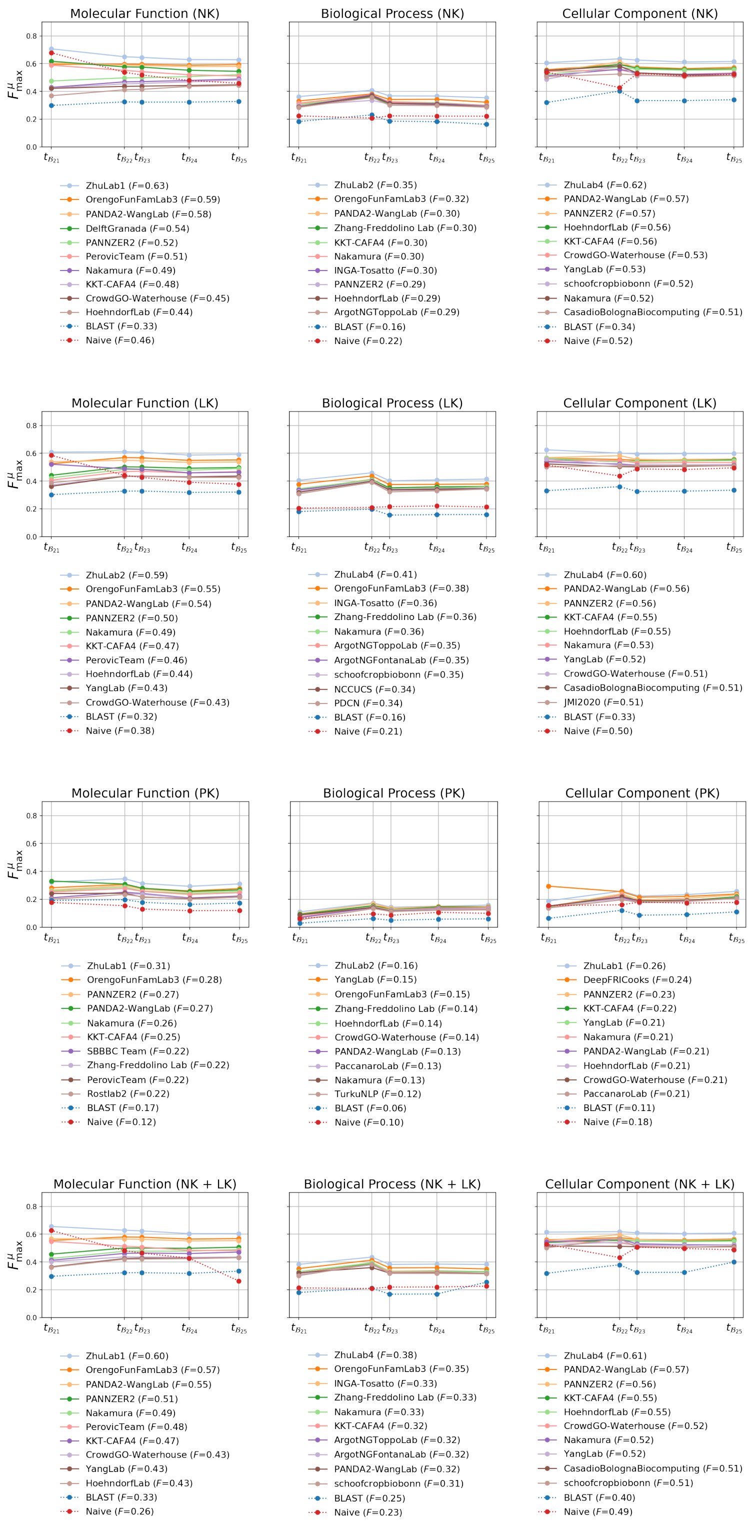


Figure S3.36: (a) $F^{\mu}_{\text{max}}$ over the years for top 10 methods for No Knowledge (NK) (b) $F^{\mu}_{\text{max}}$ over the years for top 10 methods for Limited Knowledge (LK) (b) $F^{\mu}_{\text{max}}$ over the years for top 10 methods for partial knowledge (PK) (d) $F^{\mu}_{\text{max}}$ over the years for top 10 methods for No Knowledge (NK) + Limited Knowledge (LK)

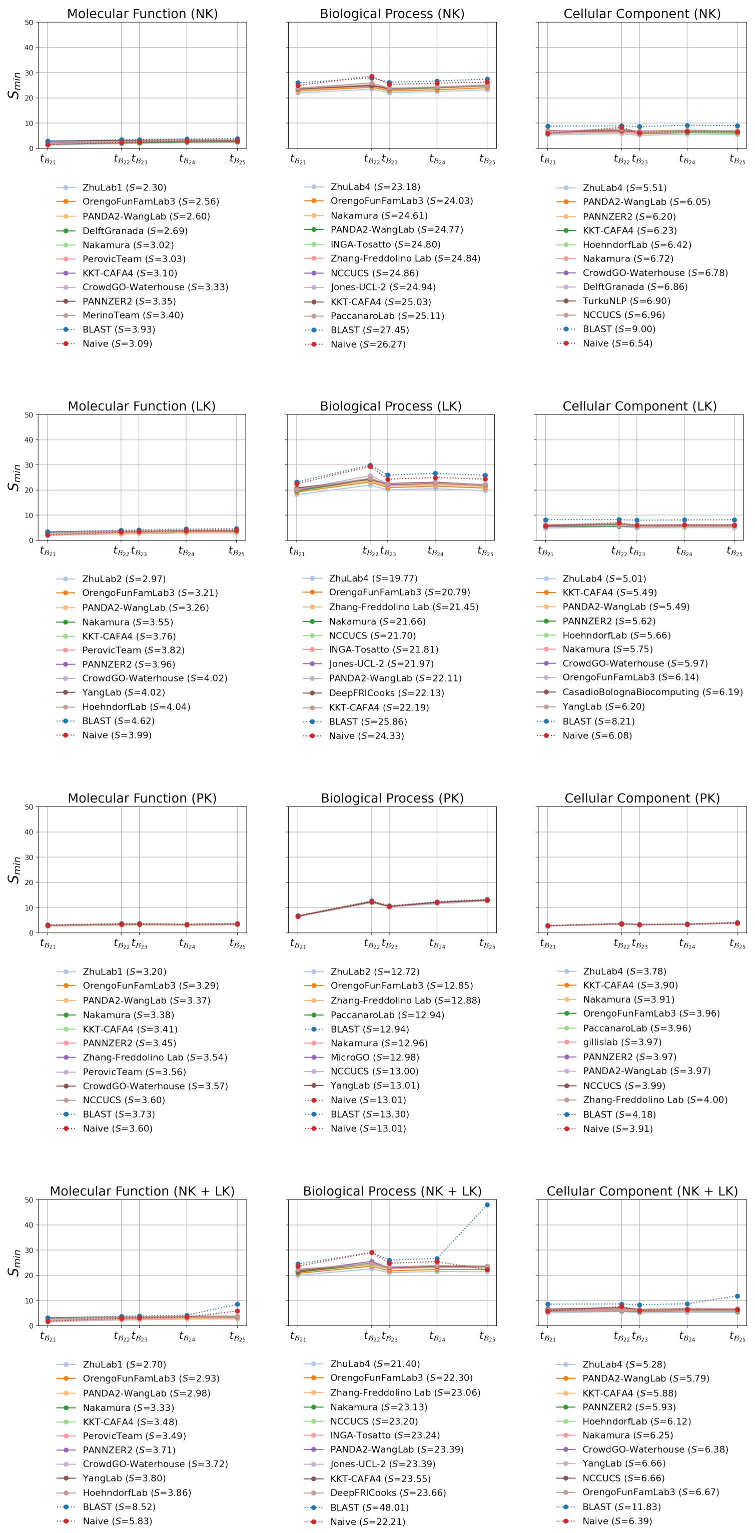


Figure S3.37: (a) $S_{\text{min}}$ over the years for top 10 methods for No Knowledge (NK) (b) $S_{\text{min}}$ over the years for top 10 methods for Limited Knowledge (LK) (b) $S_{\text{min}}$ over the years for top 10 methods for partial knowledge (PK) (d) $S_{\text{min}}$ over the years for top 10 methods for No Knowledge (NK) + Limited Knowledge (LK)
